# Extending Tests of Hardy-Weinberg Equilibrium to Structured Populations

**DOI:** 10.1101/240804

**Authors:** Wei Hao, John D. Storey

## Abstract

Testing for Hardy-Weinberg equilibrium (HWE) is an important component in almost all analyses of population genetic data. Genetic markers that violate HWE are often treated as special cases; for example, they may be flagged as possible genotyping errors or they may be investigated more closely for evolutionary signatures of interest. The presence of population structure is one reason why genetic markers may fail a test of HWE. This is problematic because almost all natural populations studied in the modern setting show some degree of structure. Therefore, it is important to be able to detect deviations from HWE for reasons other than structure. To this end, we extend statistical tests of HWE to allow for population structure, which we call a test of “structural HWE” (sHWE). Additionally, our new test allows one to automatically choose tuning parameters and identify accurate models of structure. We demonstrate our approach on several important studies, provide theoretical justification for the test, and present empirical evidence for its utility. We anticipate the proposed test will be useful in a broad range of analyses of genome-wide population genetic data.

## 1 Introduction

Hardy-Weinberg Equilibrium (HWE) is a general and far-reaching principle in population genetics that is incorporated in a wide range of applications. In evolutionary terms, HWE says that for a population meeting certain conditions the genotype frequencies of a genetic locus can be expressed in terms of the allele frequencies. The equilibrium portion of HWE comes from the fact that even if this relation does not hold in the initial state of a population, then one generation of random mating guarantees that HWE will hold. An equivalent way to frame HWE is in probabilistic terms, where the relationship between genotypes frequencies as a function of the allele frequency is a result of a Binomial distribution parameterized by the allele frequency for diallelic markers (or the Multinomial by for multiallelic markers). The genotype of a randomly sampled individual is then determined by a draw from the Binomial distribution. Tests for HWE in practice usually involve verifying the Binomial distribution of the genotypes in terms of allele frequencies.

These simple statements about HWE motivate why testing for HWE is a common preliminary step in a variety of genomic analyses — indeed, HWE serves as a data quality check or model assumptions check since it is expected to approximately hold for most markers. In studies such as genome-wide association studies (GWAS), HWE is treated as a baseline for quality control where markers deviating strongly from HWE are filtered out as likely genotyping error [1,2]. Further, the probabilistic statement of HWE where genotypes can be modeled using the Binomial distribution serves as the basis in many statistical methods. For instance, HWE serves as an assumption in models of population structure [3,4], the calculation of genetic relationship matrices [5], and in tests for genetic association [6]. Even when HWE is not explicitly stated to be an assumption, many common statistical genetic operations utilize the Binomial form of HWE, for example, scaling by the standard deviation of the Binomial in terms of allele frequencies before performing a principal components analysis or forming a genetic related matrix.

While the broad importance of HWE to genetics is clear, it is nonetheless the case that the conditions necessary for HWE are restrictive, especially in its requirement that there is no population structure present. Considering a probabilistic approach, HWE treats samples at a marker to be identical and independently distributed, i.e., completely homogenous with no structure. Population structure is ubiquitous in natural populations and therefore they are likely to violate the no structure assumption of HWE. This typically results in the appearance that the large proportions of markers deviate from HWE, obfuscating the important deviations such as those resulting from genotyping error or evolutionary selection.

These limitations are evident in how practitioners apply HWE to human genetic data. One approach is to test for HWE separately within subsets of the samples where there is less population structure. Then, test results are aggregated at each marker and some criteria accounting for the separate tests is applied to determine whether the HWE is violated. This often appears in studies where there are known population labels for the samples (for example, [7,8]). Another approach is to reject based on a very conservative p-value threshold which can vary between studies. For instance, in ref. [9], a metaanalysis study of 22 separate GWAS, the HWE p-value threshold used in each individual GWAS ranged from 10^−20^ to 10^−3^. The goal of these approaches is to reduce the number of markers violating HWE to ensure that only genotyping errors are removed.

We propose a procedure for testing for Hardy-Weinberg Equilibrium that allows for population structure, called the “structural HWE” (sHWE) test. We address these traditional limitations of HWE by extending the probabilistic model to allow for heterogeneity in the samples, i.e., by modeling the genotypes at a marker using individual-specific allele frequencies that account for structure. Individual-specific allele frequencies are the most general parameterization of structure in that there is possibility a unique allele frequency for each marker and individual combination, and common models of population structure can be formulated in this way, including the often used admixture models. We discuss specific parameterizations of this model in Section 2.3. Like current methods for testing for HWE, our proposed test of sHWE can be applied on a marker-to-marker basis to determine which markers violate HWE extended to allow for population structure. Further, the genome-wide joint distribution of sHWE p-values can be used to assess a global goodness-of-fit of the model of population structure. This allows one to choose optimal values of tuning parameters such as the latent dimensionality of a model or the number of admixed ancestral populations. Lastly, the assumptions of the sHWE test satisfy the conditions needed for association testing while controlling for population structure [10].

To illustrate the flexibility of and motivate the sHWE procedure, we analyzed single nucleotide polymorphism (SNP) genotypes from the 1000 Genomes Project (TGP) [11, 12]. The TGP data exhibits population structure in two challenging ways: first, samples were taken from populations on a global scale (including samples originally in the HapMap project), and second, some samples such as the Hispanic Latin American populations are known to have undergone recent admixture [13–15]. To model population structure, we used the logistic factor (LFA) analysis method [16], which uses *K* latent variables to account for structure. Increasing *K* captures progressively more of the population structure.

Figure 1 shows p-value histograms from this analysis, where each histogram is composed of p-values resulting from testing for HWE or sHWE on all markers simultaneously. *A* traditional test for HWE is heavily skewed towards zero, indicating that the vast majority of SNPs would be found to deviate from HWE due to the presence of population structure. Fitting the LFA model with *K* = 3 partially accounts for the population structure, and the sHWE p-values have a smaller peak at zero and less skew towards zero than the uncorrected HWE p-values. We show in Section 3.1 that *K* = 12 is the optimal value for the TGP dataset. For *K* = 12, the vast majority of genome-wide p-values are Uniform(0,1) distributed. This is the distribution of p-values when sHWE holds, and it indicates that population structure has been accounted for in our model, with the exception of the small number of SNPs found to be out of sHWE and thus also out of HWE. Since the LFA fit with *K* = 12 optimally accounts for population structure and the sHWE test incorporates population structure into the procedure, these deviations from sHWE can be attributed to technology errors or evolutionary effects.

**Figure 1:**
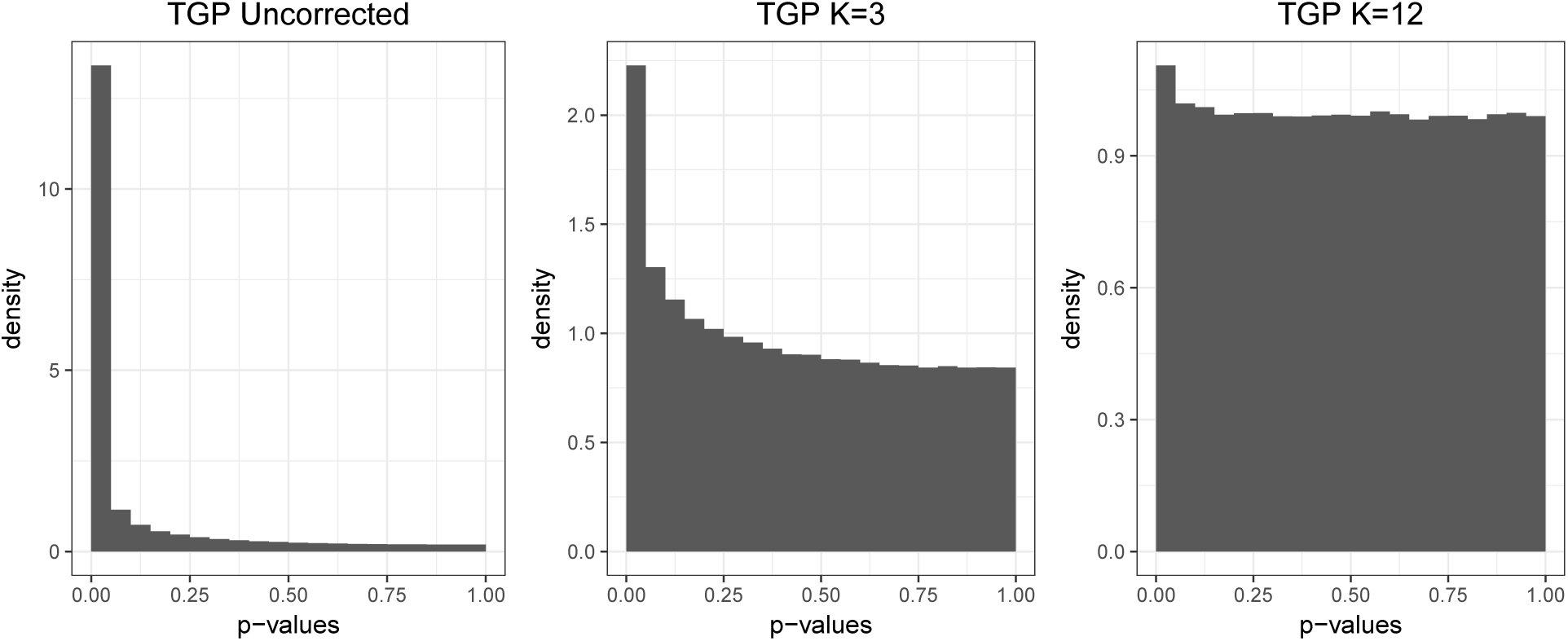
A proof-of-concept of the sHWE procedure. We fit the LFA model of structure [16] to the TGP dataset and varied *K*, which is the number of latent factors to account for population structure. The left-most panel depicts a histogram of genome-wide p-values for a traditional test of HWE, which is equivalent to using the sHWE test with a population structure model of dimensionality *K* = 1. The histogram is heavily skewed towards zero, showing that most SNPs would be identified as deviating from HWE. The middle panel depicts sHWE test p-values for *K* = 3, which partially accounts for the population structure. As a result, there is less skew towards zero, and the large p-values (i.e., larger than 0.75) are Uniform distributed which indicates that some SNPs are in sHWE. The right-most panel depicts sHWE test p-values for *K* = 12, the empirically optimal value, which best accounts for population structure in the dataset. The SNPs concentrated at the peak near zero are found to be deviated from sHWE, indicating that they violate HWE for reasons other than population structure.

The sHWE test is performed by fitting a model of population structure that parameterizes allele frequencies for each individual and single nucleotide polymorphism (SNP) pair. Then, we simulate null genotyping datasets that preserve the observed population structure where sHWE holds. Finally, we calculate a test statistic that measures deviation from sHWE for the observed and null datasets, and a p-value is computed. The algorithm is shown in Figure 2. We demonstrate our method on several publicly available global human datasets: the Human Genome Diversity Project (HGDP) [17], the 1000 Genomes Project [11,12], and a dataset genotyped using the Affymetrix Human Origins chip [18]. We first analyze these datasets independently, showing how the sHWE procedure allows one to choose the dimensionality of the population structure model. Then, we compared SNPs that are misspecified with respect to the population structure model between datasets and technologies, showing that the sHWE procedure identifies SNPs affected by genotyping errors and that results are replicable between datasets.

**Figure 2:**
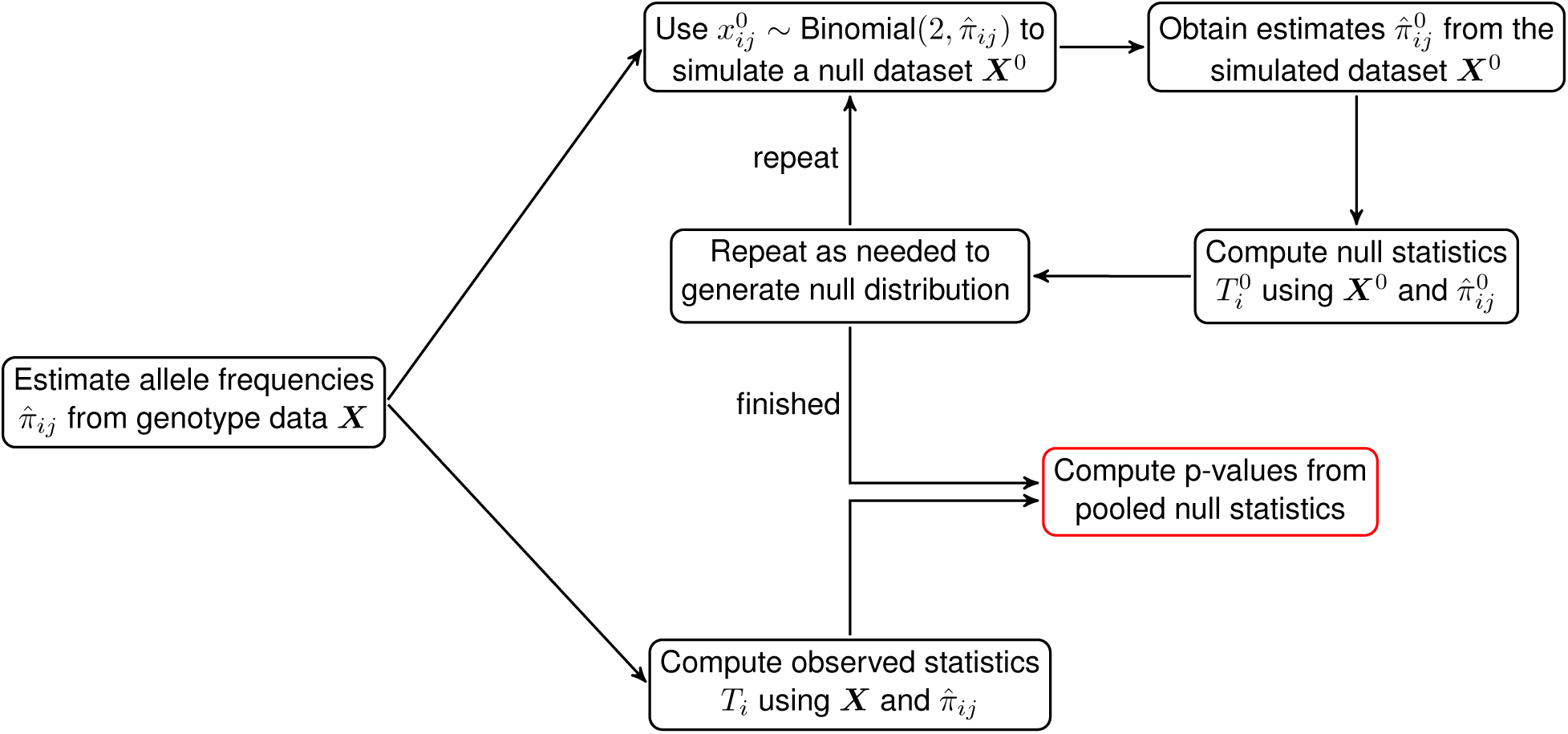
The structural HWE testing procedure as a schematic. Using the genotype matrix ***X***, we first fit a model of population structure to estimate 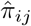. The values of 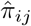 are used to simulate null datasets incorporating the sHWE assumptions. We compute sHWE test statistics for both observed and simulated null datasets and compute p-values by comparing the values of the observed test statistics and the pooled null test statistics.

## 2 Methods

We first introduce the globally sampled human genome-wide genotyping datasets used in this analysis. Then, we show how the probabilistic interpretation of HWE can be extended to the sHWE test by incorporating the most general representation of population structure. We discuss a few ways to parameterize population structure and consider how the sHWE test behaves when all parameters are known. In addition, we show how to implement the sHWE test in practice using simulated empirical null distributions based on genome-wide genotyping data. Finally, we discuss how the sHWE procedure can be used to validate and tune models of population structure in addition to the standard application of marker quality control.

### 2.1 Datasets

We used genome-wide genotyping data from three publicly available sources, each of which performs global sampling of humans.

#### Human Genome Diversity Project (HGDP)

This study sampled globally from 51 populations [17]. We filter for related individuals using the “H952” subset from ref. [19]. Genotypes were filtered with minimum allele frequency of 0.05 and minimum genotype completeness of 0.995. The dimensions of this dataset are 940 individuals and 550,303 SNPs. The data are available at http://www.hagsc.org/hgdp/files.html.

#### Human Origins (HO)

This study sampled globally from 147 populations [18]. These samples were genotyped on the Affymetrix Human Origins array, which was specially designed for population genetics applications. We used the publicly available portion of the dataset. Genotypes were filtered with minimum allele frequency of 0.05 and minimum genotype completeness of 0.99. After filtering the dataset for ancient and non-human samples, we are left with 372446 SNPs and 2,251 individuals. The data are available from the Reich lab webpage: http://genetics.med.harvard.edu/reich/Reich_Lab/Datasets.html.

#### 1000 Genomes Project (TGP)

This study analyzed genome sequence diversity in humans through whole genome sequencing [11,12]. They also provide SNP chip genotyping on the Illumina Omni platform for the Phase 3 release. Genotypes were filtered with minimum allele frequency of 0.05 and minimum genotype completeness of 0.99. After removing related individuals, the dataset consists of 1,815 individuals and 1,229,310 SNPs. The data are available at ftp://ftp.1000genomes.ebi.ac.uk/vol1/ftp/release/20130502/supporting/hd_genotype_chip/.

Further, we generated two additional datasets from TGP for the purpose of comparing individuals genotyped on different technologies: one from variants called from sequencing data (the TGP phase 3 variant calls available at ftp://ftp.1000genomes.ebi.ac.uk/vol1/ftp/release/20130502/) and the other a version of the genotyping chip data described above. Both datasets were designed to maximally overlap. We filtered both datasets to share the same unrelated individuals (1,683 individuals total). We used a subset of 1,224,056 of the SNPs from the genotyping chip data overlapping with the variant calls. The variant calls were designed to include as many SNPs from the chip data as possible, as well as additional SNPs that were least 5kbp. This resulted in a subset of 1,306,465 SNPs from the variant calls.

### 2.2 Traditional HWE as a probability model

Typically, population genetic assumptions such as infinite population size, random mating, no selection, no mutation, and no migration (among others) are assumed as the starting point for HWE. At a particular locus with alleles *A* and *B* and allele frequencies *p* (corresponding to allele *A*) and *q* = 1 − *p* (corresponding to allele *B*), HWE states that after one generation of random mating, the genotype frequencies of *AA, AB*, and *BB* are *p*^2^, 2*pq*, and *q*^2^, respectively. The allele frequencies and genotype frequencies then remain at these values for all further generations. HWE can be viewed as a probabilistic model if we consider *A* to be the reference allele and code the genotypes as 0, 1, and 2, corresponding to *BB, AB*, and *AA*, respectively. The genotype for each individual at this locus is modeled under HWE as an independent draw from a Binomial(2, *p*) distribution. The relationship between genotype frequency and allele frequency follows directly from this distributional assumption.

Common tests for HWE such as Pearson χ^2^ test for goodness-of-fit or Fisher’s exact test [20] test for whether the observed genotype counts are compatible with draws from a Binomial distribution using the observed allele frequency as the probability of success. Many of the population genetics assumptions are related to the statistical assumption that alleles and individuals can be treated as independent and identically distributed. In human datasets, the presence of population structure means this assumption is violated, resulting in a deviation from HWE. We will directly account for population structure by parameterizing it in a way that is compatible with a Binomial model of genotypes so that this probabilistic model of HWE can be tested in the presence of structure.

### 2.3 Models of population structure

Consider a dataset consisting of *m* diallelic SNP genotypes (coded as 0,1, and 2 copies of the reference allele) and *n* individuals. For current GWAS, *m* is often on the order of millions while *n* is in the tens of thousands. We will aggregate the data into a genotype matrix *X* with dimensions *m* by *n*, and choose indices such that *x_ij_* corresponds to the *i*-th SNP and *j*-th individual in *X*.

For complete generality, let us allow each individual and SNP pair to have its own reference allele frequency *π_ij_*. We can aggregate the *π_ij_* into a *m × n* matrix *F* whose (*i, j*) element is *π_ij_* and 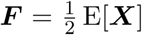. This represents the most general way to probabilistically represent population structure, as there are as many parameters as there are individual-SNP pairs. Models of population structure typically parameterize *π_ij_* with constraints, such that fewer parameters are needed. Using more sophisticated parameterizations of *π_ij_* will allow us to relax the statistical assumption that the individuals are all identically distributed.

We will now summarize several special cases of the above general parameterization, where there are constraints on the *π_ij_* values. In all cases, these models assume that the genotypes are generated independently according to *x_ij_* ~ Binomial(2, *π_ij_*). The simplest parameterization of *π_ij_* is that of a population in HWE with no structure, where all *π_ij_* = *p_i_* and *p_i_* is the observed allele frequency at SNP*i*. In a model with non-overlapping, independently evolving subpopulations, there are *K* subpopulations and *π_ij_* is the allele frequency of SNP *i* for the subpopulation of which individual *j* is a member. In an admixture model of population structure [3, 21], there are *K* ancestral populations, and the relevant model parameters are *q_j_*, the *K*-vector of admixture proportions for individual *j*, and *p_i_*, the *K*-vector of allele frequencies for SNP *i*. Then, *π_ij_* is the weighted sum of these parameters, 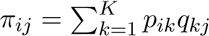. In spatial models of population structure [22,23], *π_ij_* is explicitly a smooth function of the geographical coordinates of each sample.

In this paper, we focus on two approaches from ref. [16]: the logistic factor analysis (LFA) and the truncated principal components analysis (PCA). These methods are both computationally efficient and were shown to outperform existing methods for estimating ***F***. They directly model *π_ij_* using low-dimensional factorizations and are accurate and computational efficient on large datasets [16]. We primarily use the LFA method, which models *π_ij_* using its natural parameterization, logit(*π_ij_*) = log (*π_ij_*/(1 − *π_ij_*)). Population structure is captured by factorizing the logit transformation of ***F***: logit(***F***) = ***AH***, where ***A*** is a *m* by *K* matrix and ***H*** is a *K* by *n* matrix. The columns of ***H*** represent population structure for each individual, while the rows of ***A*** are the way population structure is manifested in each SNP. We also show results for the PCA approach to estimating *π_ij_*.

The truncated PCA approach utilizes the fact that under the binomial model relating *x_ij_* and *π_ij_*, E[*x_ij_*] = 2*π_ij_*. The estimates of *π_ij_* are formed by projecting ***X*** onto first *K* principal components of ***X*** and scaling the projection by a factor of 1/2. Some of the values for 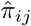 may be outside of the interval [0,1], in which case they are replaced by 1/(2*n*) or 1 − 1/(2*n*), which corresponds to the allele frequency for having only one copy of an allele. In addition to these two methods, we also demonstrate our proposed test on the ADMIXTURE [21] method for modeling population structure through the probabilistic admixture model described above.

All of the models of population structure summarized here involved a tuning parameter, such as the number of ancestral populations *K*, a smoothing parameter in the spatial model, or the number of latent factors *K*. Our proposed method will introduce a way to automatically choose the value of these tuning parameters by considering the goodness of fit of the model of structure to the data.

### 2.4 A structural test for Hardy-Weinberg equilibrium

Using the individual-specific allele frequencies *π_ij_* offers a general framework for extending tests of HWE to allow for structure. To this end, we will derive a test for “structural Hardy-Weinberg equilibrium” (sHWE) by extending the derivation of the standard Pearson χ^2^ statistic.

The data for a single marker can be summarized using the genotype counts for a SNP, which we will call *N*^(^*^G^*^)^, where *G* ∈ {0,1,2} are the different genotypes. We put the genotype variable in parentheses and as a superscript to distinguish it from the indices for matrices and vectors. We will introduce a new test statistic that is to be calculated for each SNP. Thus, consider a fixed SNP and drop the corresponding subscript *i*. This leaves us with the vector of genotypes *x* = (*x*_1_, *x*_2_,…, *x_n_*) and a vector of allele frequencies *π* = (*π*_1_, *π*_2_,…, *π_n_*). The test of sHWE performs a hypothesis test on the distribution of the genotype data as follows:

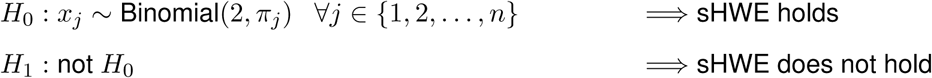

This hypothesis test is performed for all markers simultaneously, with the goal of identifying which markers deviate from the sHWE assumptions.

We can write the genotype counts as 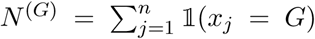. We can define the quantity 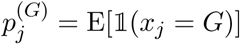, which depends on the genotype *G* in the following way when sHWE holds:

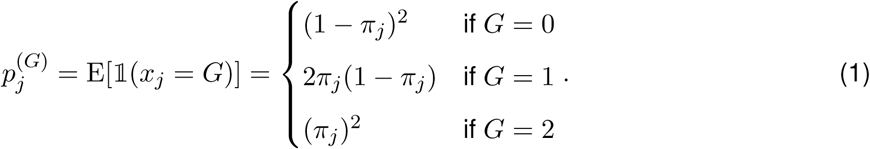

This notation will allow us to consider the distribution of *N*^(^*^G^*^)^ and to formulate a test where the null hypothesis is that Equation (1) holds for all *j* and the alternative is that Equation (1) does not hold for at least one *j*. It follows that 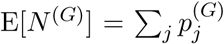 and 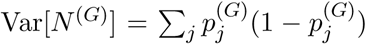. We can apply the central limit theorem to note that *N*^(^*^G^*^)^ is asymptotically distributed as a Normal with mean 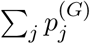 and variance 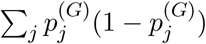.

Now consider just two of the genotype counts as a vector of length two called *v* = (*N*^(0)^, *N*^(1)^), since *N*^(2)^ = *n*-*N*^(0)^ − *N*^(1)^. It is distributed bivariate Normal with mean vector 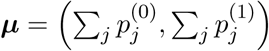 and covariance matrix Σ, where:

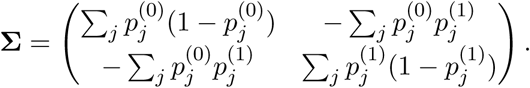

Thus, the quadratic form

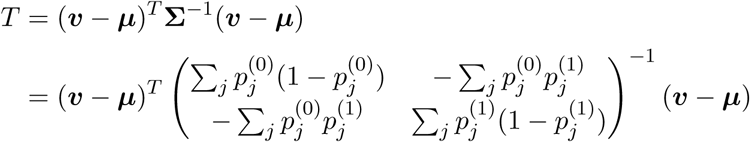
has asymptotic distribution χ^2^ with 2 degrees of freedom. We can use *T* as a test statistic for sHWE for each SNP when each *π* is known — and therefore *F* is known. We next show how to incorporate unknown ***F*** into this statistic and determine its null distribution. Note that when the null is true for unstructured HWE, the sums in the expression for Σ vanish, and *T* simplifies to the usual Pearson χ^2^ statistic, so our proposed statistic is a generalization of the usual χ^2^ test of HWE.

### 2.5 Algorithm

In practice, the *π_ij_* values are unknown and the act of estimating them from the data being tested changes the null distribution of *T* [24]. We propose computing an empirical null distribution via the parametric bootstrap [25] by simulating data from the model of population structure, computing sHWE statistics for the simulated dataset, and treating those statistic as samples from a null distribution. There are two aspects of the sHWE statistic that make the simulation of an empirical null attractive. First, models of population structure seek to account for the dependence present in data. Simulating an empirical null allows us to compute the sHWE statistic for data where the observed structure is preserved. Second, the statistic derived earlier is a pivotal quantity, i.e., the null distribution is always *χ*^2^ with 2 degrees of freedom regardless of the values of *π_ij_*.

Strictly speaking, each simulated dataset serves as one bootstrap sample for each SNP. It would be too computationally intensive to simulate the datasets needed to have enough resolution to compute meaningful p-values. However, since the sHWE statistic is a pivotal quantity, pooling the simulated null datasets is an effective strategy. The pooling procedure is to simulate a small number of null datasets, then combine the sHWE statistics for all of the simulated SNPs as observations from the null distribution. We validate the use of a pooled empirical null by comparing the distribution of p-values computed using the pooling routine and the distribution of marginal p-values, where an empirical null distribution is calculated for each SNP. We show quantile-quantile plots in Figure S1 between the p-values computed using the two types of null distributions. Their joint distributions are nearly identical, thus validating the pooled empirical null procedure [26].

Our algorithm to test for sHWE in a dataset of SNPs is described in Algorithm 1. A graphical depiction of the algorithm is shown in Figure 2.

### 2.6 Structural HWE as empirical model tuning and validation

Since many models of population structure parameterize *π_ij_*, sHWE provides a framework for validating models of population structure that parametrize *π_ij_*. By testing each individual SNP for violation of a model’s assumptions, we can aggregate the tests to determine if the overall population structure model accounts for the variation in the data appropriately. When the model is well formulated, the vast majority of SNPs should pass the sHWE test. Thus, we can examine the joint distribution of the p-values computed at every SNP in the dataset. The expected behavior of the distribution of p-values is that the they are Uniform distributed across the interval [0,1], except near zero where a small portion of SNPs are shown to deviate from sHWE by having significant p-values (e.g., Figure 3). Choosing the significance threshold can be done with a variety methods, such as false discovery rates [28]. This provides a natural criterion for filtering SNPs that violate the model assumptions, and this an important part of any robust preliminary analysis.

**Algorithm 1.**
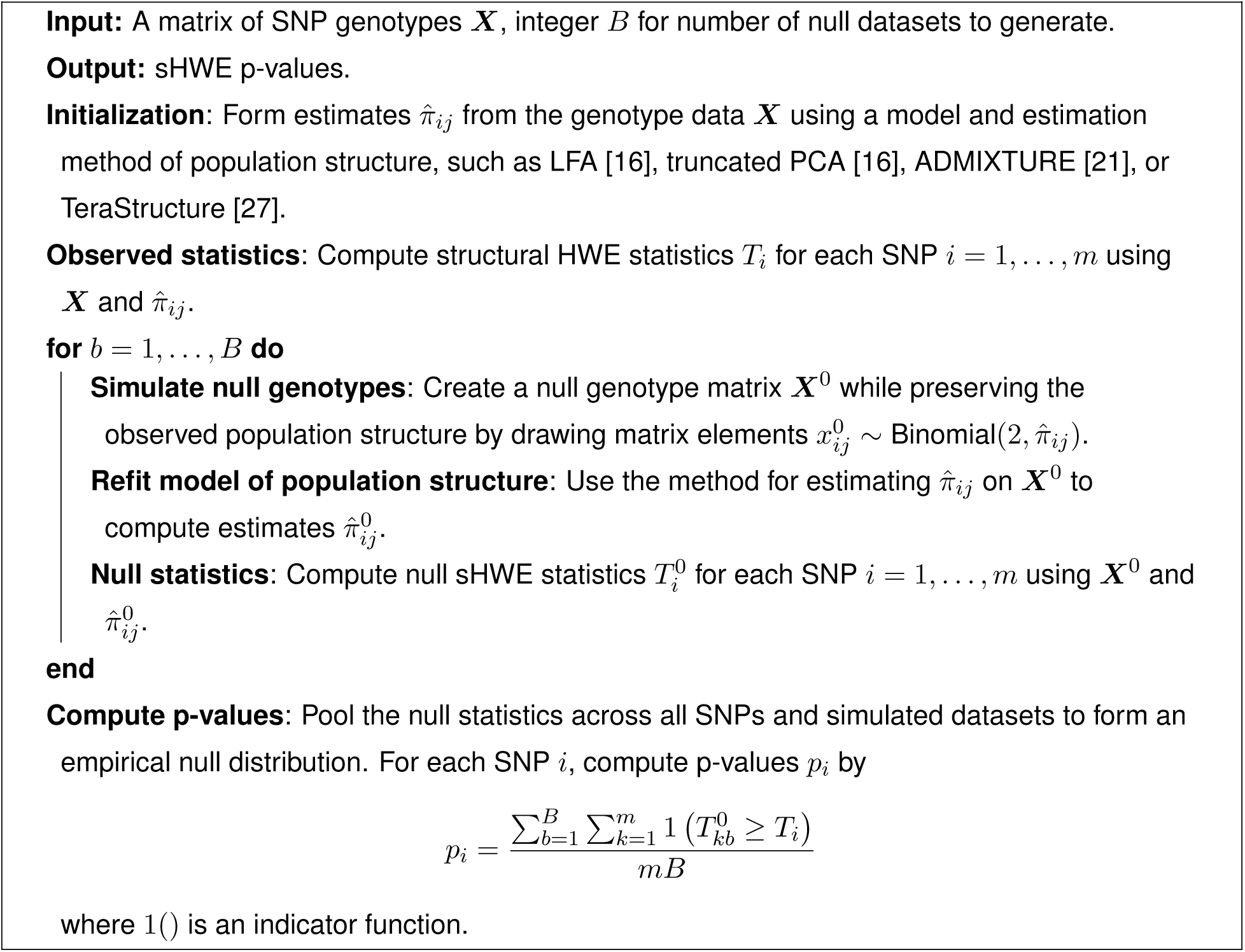
Procedure for computing genome-wide sHWE p-values.

**Figure 3:**
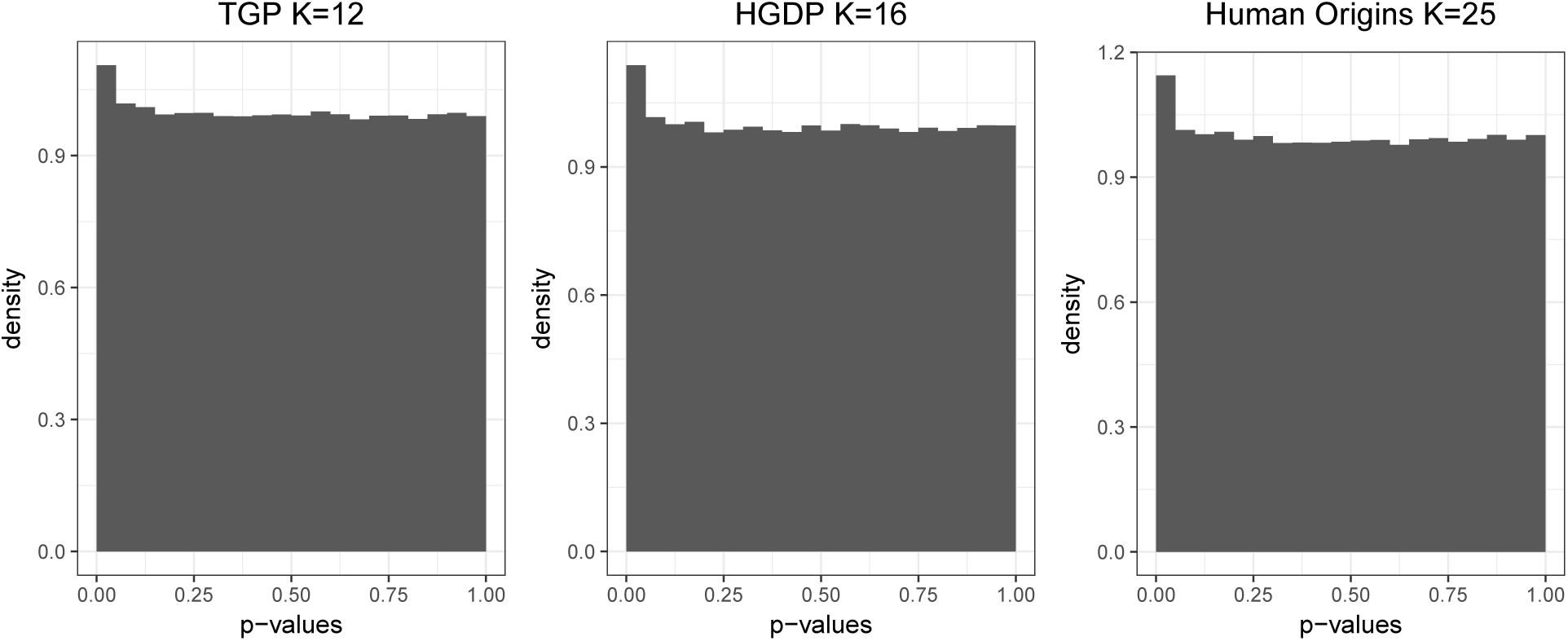
Histogram of sHWE test p-values for each dataset at chosen *K* as determined by the entropy measure. The sHWE test is performed for each SNP in the dataset after fitting the LFA model of population structure. The aggregated p-values are mostly Uniform(0,1) distributed, except for a peak at 0. This indicates that most of the SNPs are in sHWE, given the fitted structure. The peak at 0 contains an enrichment of SNPs that deviate from sHWE.

This leads to a principled procedure for optimizing tuning parameters in the model of population structure such as the latent dimensionality *K*. If we compute sHWE p-values for a range of *K*, we can choose the value of *K* that has the best null properties. It is important to distinguish the characteristic of having good null properties from an absolute measure like least number of significant SNPs. Because our procedure is verifying a model fit over the genome, we want to choose the parametrization of nowhere the p-values are most Uniform over the largest possible interval, excluding a possible peak near 0. The algorithm is detailed in Algorithm 2. Note that while we found to *C* = 150 to be sufficient for analyses with 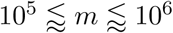, it may be helpful to choose a higher value if there are many more SNPs (or lower value for smaller datasets).

**Algorithm 2.**
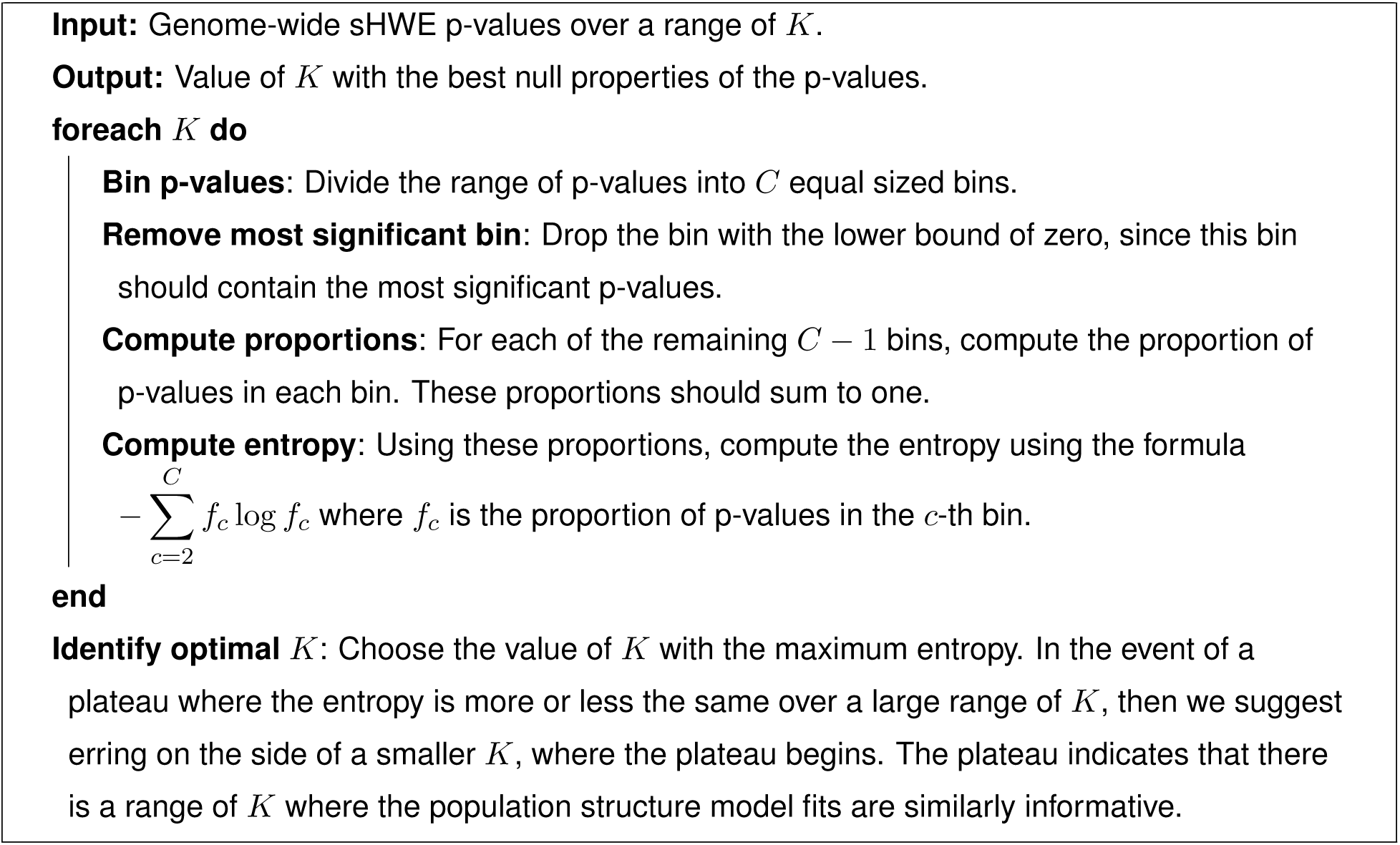
Entropy based procedure for automatically choosing the value of *K*.

### 2.7 Software

Our procedure is implemented in the `ifa` R package [16] (available at http://github.com/StoreyLab/ifa) as the function `sHWE`.

## 3 Results

We demonstrate the sHWE procedure on the global datasets detailed in Section 2.1. We also show that the sHWE procedure works with the truncated PCA method [16] and the ADMIXTURE method of fitting population structure [21]. Then, we consider a few ways to interpret the results of testing for sHWE. First, we show that there are no systematic differences in sHWE p-values when SNPs are separated by annotations or minor allele frequency. Then, we consider the replicability of sHWE results between the global datasets, as well as differences between results for the TGP samples on two different genotyping technologies.

### 3.1 Analyzing global datasets

We demonstrate our proposed procedure where *π_ij_* is estimated using the LFA method [16] on three highly structured and global datasets: the HGDP, *HO*, and TGP (genotyping chip) datasets described in Section 2.1. We utilized *B* = 3 null simulations from Algorithm 1 in the calculations. We show the sHWE p-value distributions over a range of *K*, the latent dimensionality of the LFA model of population structure, for the three datasets in Figures S2, S3, and S4. The distributions of p-values share the same general behavior between datasets. When *K* is too small and the population structure is insufficiently modeled, the sHWE test p-values are skewed heavily towards 0. As additional latent factors are added to account for more structure across the genome, the p-value distributions shift away from zero and become more uniform. Eventually, the p-value distributions become skewed towards 1, as population structure model is overfit to the data.

For each dataset, we observe that there is a range of *K* where we observe the desired distribution of p-values, i.e., a peak near zero and uniform elsewhere. Model fits in this range of *K* have the highest confidence of being well formulated and all serve equally well as a basis for future analysis. We suggest choosing *K* following the entropy measure presented in Section 2.6. We show results in Figure 4, corresponding to *K* = 12 for TGP, *K* = 16 for HGDP, and *K* = 25 for Human Origins. At these values of *K*, we estimate the proportion of SNPs that are in sHWE, using the bootstrap method from the `qvalue` *R* package [28]. We find these estimated proportion to be 0.990 for TGP, 0.990 for HGDP, and *π*_0_ = 0.989 for HO; this suggests that the vast majority of human SNPs are in HWE. SNPs in sHWE are also interpreted to be well parameterized by the LFA population structure model.

To demonstrate results using other parameterizations of *π_ij_*, we first analyze these three datasets for a range of *K* using the truncated PCA method [16]. The resulting histograms (Figures S5, S6, and S7) are comparable to those estimated using *π_ij_*, except that there is a small peak near 1 for larger values of *K*. This likely reflects noise introduced by the truncation procedure.

**Figure 4:**
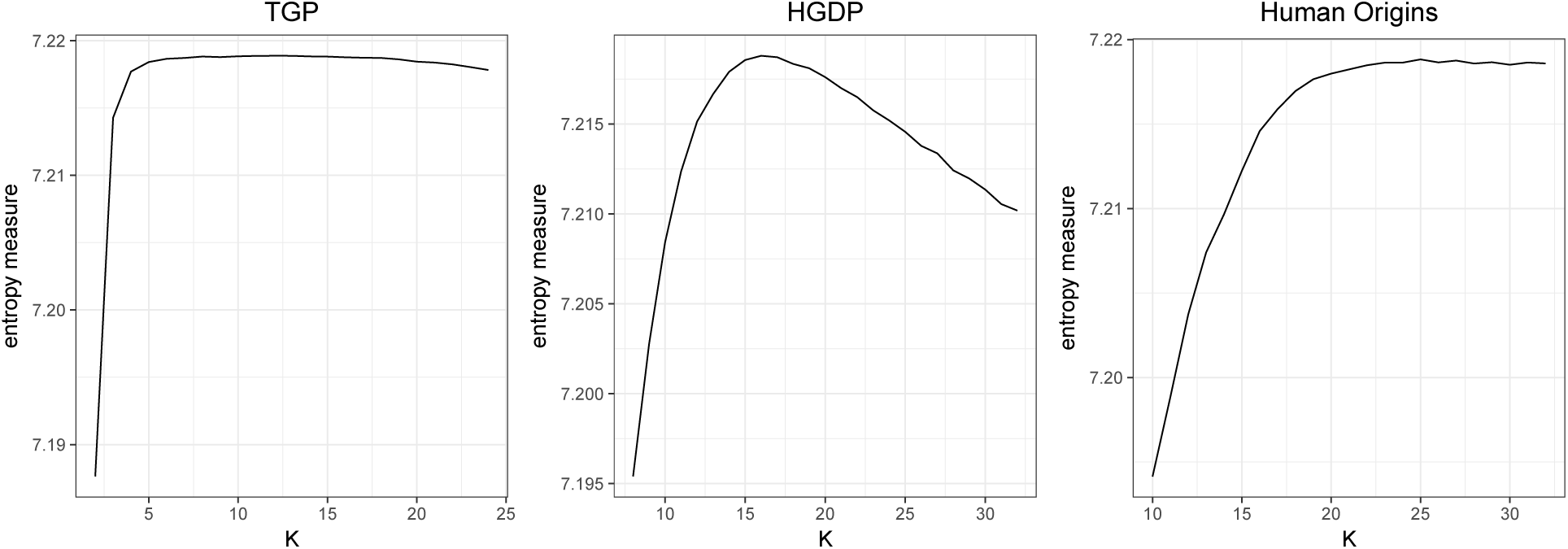
The entropy measure of uniformity of p-values for each dataset as a function of *K*. For each model fit and value of *K*, the p-values for each dataset were summarized by counting the number in each of 150 equal sized bins in the range [0,1]. The bin closest to zero was dropped, as the most significant p-values will be in that bin. The proportion of counts in the 149 bins remaining are used to compute the entropy corresponding to *K*. Higher entropy means more uniform.

We also tested parametric models of population structure. We used the ADMIXTURE method [21], which is a widely used software for fitting the admixture model of population structure, before applying the sHWE procedure. The resulting figures for the Human Origins dataset are shown in Figures S8 and S9. The sHWE p-values exhibit the expected behavior in terms of histogram shape over the range of *K*. We note that the entropy measure plateaus at *K* = 30, which is a higher value of *K* than with the LFA method on this dataset. This is expected behavior, as the admixture model is more constrained than the LFA model, since the factors need to be valid probabilities. Thus, a higher *K* is needed to achieve a similar fidelity in the modeled population structure. Further, while the sHWE procedure works with ADMIXTURE, it is a more arduous task computationally, since the individual model fits are slower than with LFA, and the sHWE procedure requires multiple fits per value of *K*.

### 3.2 The role of SNP annotation in deviations from sHWE

We compared the distributions of sHWE p-values in each dataset when separated by functional annotations of the SNPs [29]. We considered three nested levels of labels. First, SNPs were separated into intragenic and intergenic categories. Then, the intergenic SNPs were separated by whether they were in an exon or an intron. Lastly, the exonic SNPs were separated into synonymous and non-synonymous mutations. For each dataset, we found no differences between the distributions of sHWE p-values for each of the categories (Figures S10, S11, S12). Further, we found minimal differences in the distribution of p-values when binned by minor allele frequency (Figures S13, S14, S15).

### 3.3 Replicability of sHWE between datasets

To demonstrate the robustness of our sHWE procedure, we compared the results between datasets by analyzing the overlapping SNPs. For each pair of datasets, we first identified the SNPs shared by the two datasets. Between HGDP and TGP there were 357,314 shared SNPs, between Human Origins and HGDP there were 130,572 shared SNPs, and between TGP and Human Origins there were 163,443 shared SNPs. Within each of the three pairs of datasets, we compared the two sets of shared p-values by examining the most significant tail of the distribution of p-values. We chose the length of the tail by identifying how many SNPs are significant within each of the six sets of p-values at a 20% FDR threshold using the `qvalue` R package [28]. Then, for each pair, we chose the larger number of significant SNPs. The goal of this approach was to choose enough SNPs such that we capture a reasonable number of significant and non-significant SNPs. We observed concordance between the data sets because SNPs that were significant in one data set showed sHWE p-values in the other data set that are skewed towards zero and stochastically less than the Uniform(0,1) distribution. If there were no concordance we would expect these replication p-values to be approximately Uniform(0,1), which they are not. This suggests that deviations from sHWE show concordance between datasets, which in turn suggests that some of the effects driving the violation of sHWE are shared between datasets (i.e., biological) and not unique to a dataset (i.e., genotyping errors). The similarity between datasets is strongest in the comparison between the HGDP and HO datasets, which also share many of the same individuals, albeit genotyped on different technologies. These comparisons are shown in Figure 5.

**Figure 5:**
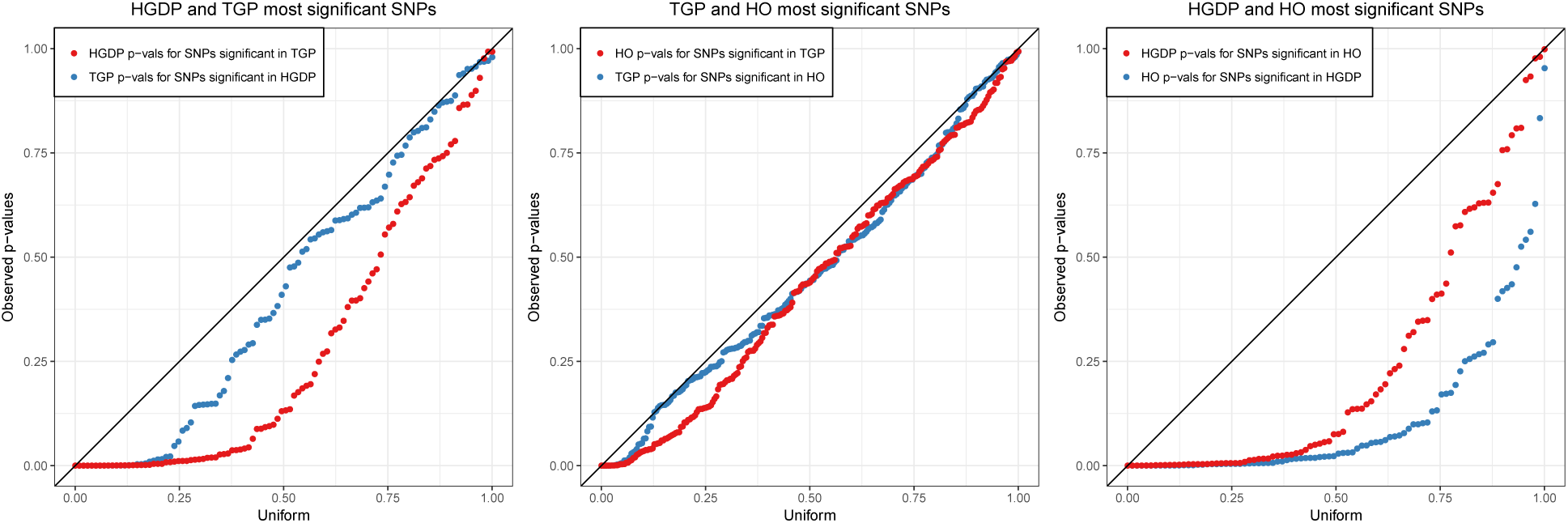
Comparisons of significant sHWE p-values between the three datasets. For each pair of datasets, we choose the *S* most significant SNPs from one dataset, where *S* is the greater of the number of significant SNPs at FDR q-value ≤ 20% for both datasets. We then test the corresponding *S* SNPs for sHWE in the other data set. Quantile-quantile plots of the resulting p-values versus the Uniform(0,1) quantiles shows that the deviations from sHWE are enriched in the other data set, verifying concordance of departures from sHWE between data sets.

### 3.4 sHWE between genotyping technologies

The 1000 Genomes Project provides a controlled setting to further investigate sHWE in genome-wide data because samples have been genotyped using different technologies. In addition to genotyping chip data, we also incorporate the integrated variant callset made by the 1000 Genomes Project, which are derived primarily from sequencing data. We created a subset of both the genotype chip data and integrated callset that share the exact same individuals while maintaining a maximal overlap in the SNPs (see Methods). We then calculated sHWE p-values for both datasets at *K* = 12, which was determined earlier for the TGP dataset.

To compare the results between the technologies, we employed two approaches. First, we analyzed the shared SNPs between the datasets generated using an approach identical to the between datasets comparison earlier, using maximum number of significant SNPs at FDR q-value ≤ 20% (Figure 6). We observe that the majority of sHWE p-values in one dataset for SNPs that are significant in the other are extremely small, though they are not necessarily significant at this particular significance threshold. This still represents concordance as the p-values in the other dataset are stochastically much smaller than Uniform. The remaining p-values are linear, meaning that the right tail of the p-values are approximately uniform away from 0. This shows that the majority of SNPs (approximately 75-80%) that deviated from sHWE exhibit this behavior in both datasets.

**Figure 6:**
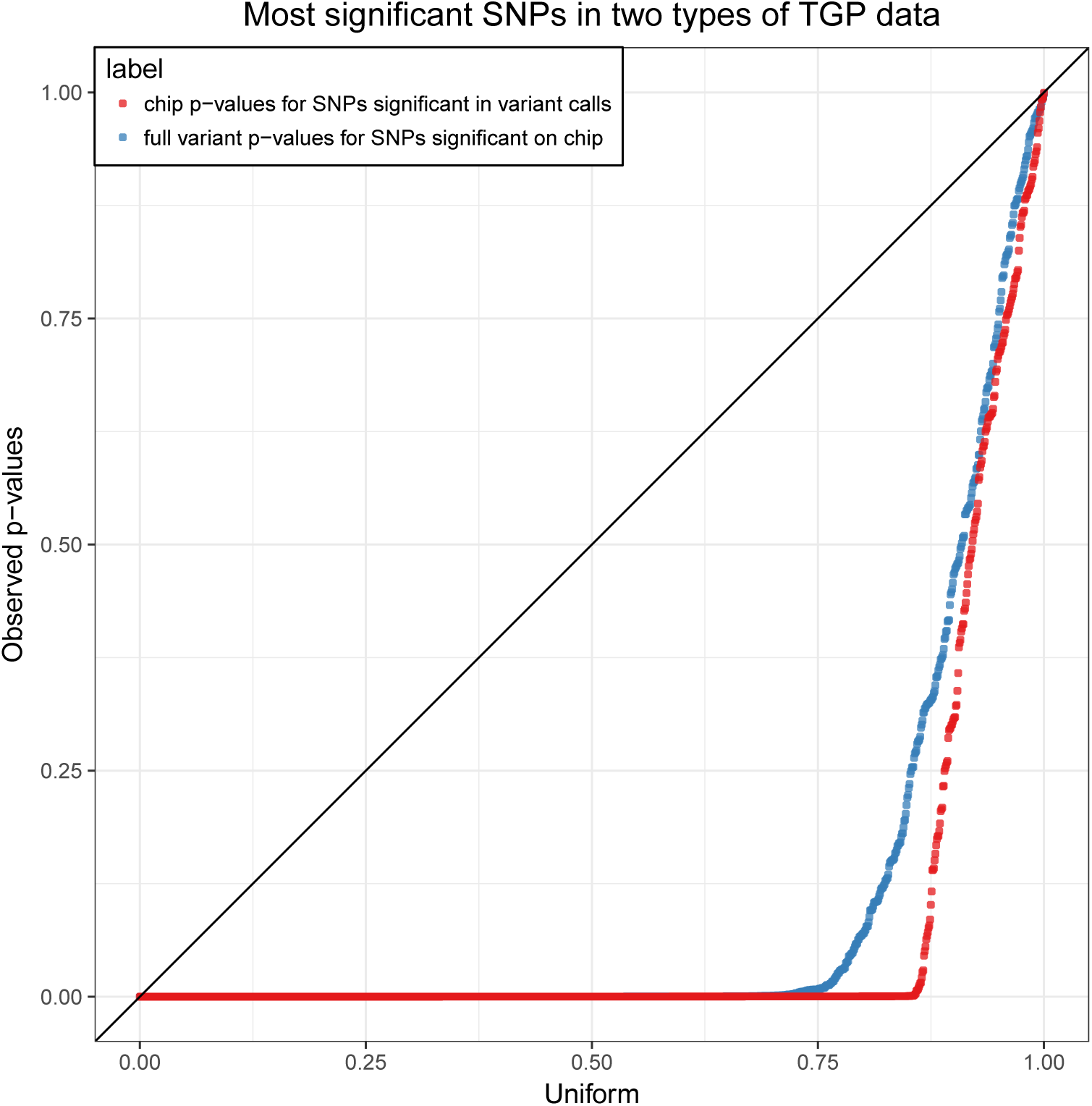
Comparisons of sHWE p-values between TGP genotyping array data and the TGP variant calls. We identify significant SNPs at FDR q-value ≤ 20% for the two datasets, then plot quantile-quantile plots of the SNPs shared in the other dataset against the Uniform distribution. The labelings follow the convention in Figure 5.

Then, we compared the distributions of all sHWE p-values between the two datasets for the shared and unshared SNPs (Figure S16). While the distribution of sHWE p-values for all shared SNPs is nearly identical, there are proportionally many more significant SNPs deviated from HWE in the integrated variant callset than in the genotyping chip data. This suggests that SNPs called from the sequencing data are less accurate than those genotyped using chips.

## 4 Discussion

We extended the Pearson χ^2^ test of HWE to allow for population structure, called the structural HWE (sHWE) test. This allows one to identify genetic markers that deviate from HWE for reasons other than population structure. For example, SNP markers can be identified in a GWAS with structure that potentially have genotyping errors, or genetic loci that are subject to evolutionary forces of interest other than structure can be identified for further analysis.

Our proposed approach is flexible in terms of the exact formulation of the model of structure. It only requires that each SNP and individual pair is drawn from a Binomial distribution, which is a condition satisfied by most common models of population structure. We chose to employ the logistic factor analysis model here [16], which serves as a base model of population structure for a test of association in GWAS [10]. A caveat of our sHWE procedure is that simulating the empirical null distribution means that we are reliant on computationally efficient methods for modeling population structure.

We demonstrated the proposed sHWE test on three highly structured global datasets. In each dataset, we showed there is a configuration of the population structure model that captures the full range of genetic variation for ~ 99% of the SNPs and that the testing procedure provides a metric for choosing the dimension.

Model-validation is an important preliminary step when applying probabilistic models to genome-wide genotyping data. We have shown that our sHWE test is a powerful tool for doing so. This approach to goodness-of-fit is applicable to any high-dimensional latent structure model for which it is possible to efficiently simulate data from a given model fit. Further, our sHWE procedure yields the ability to examine a wider range of biological questions, as our understanding of deviations from HWE in unstructured populations can now be applied to structured populations.

## Acknowledgements

This research was supported in part by NIH grant HG006448.

## 5 Supplementary Figures

**Figure S1:**
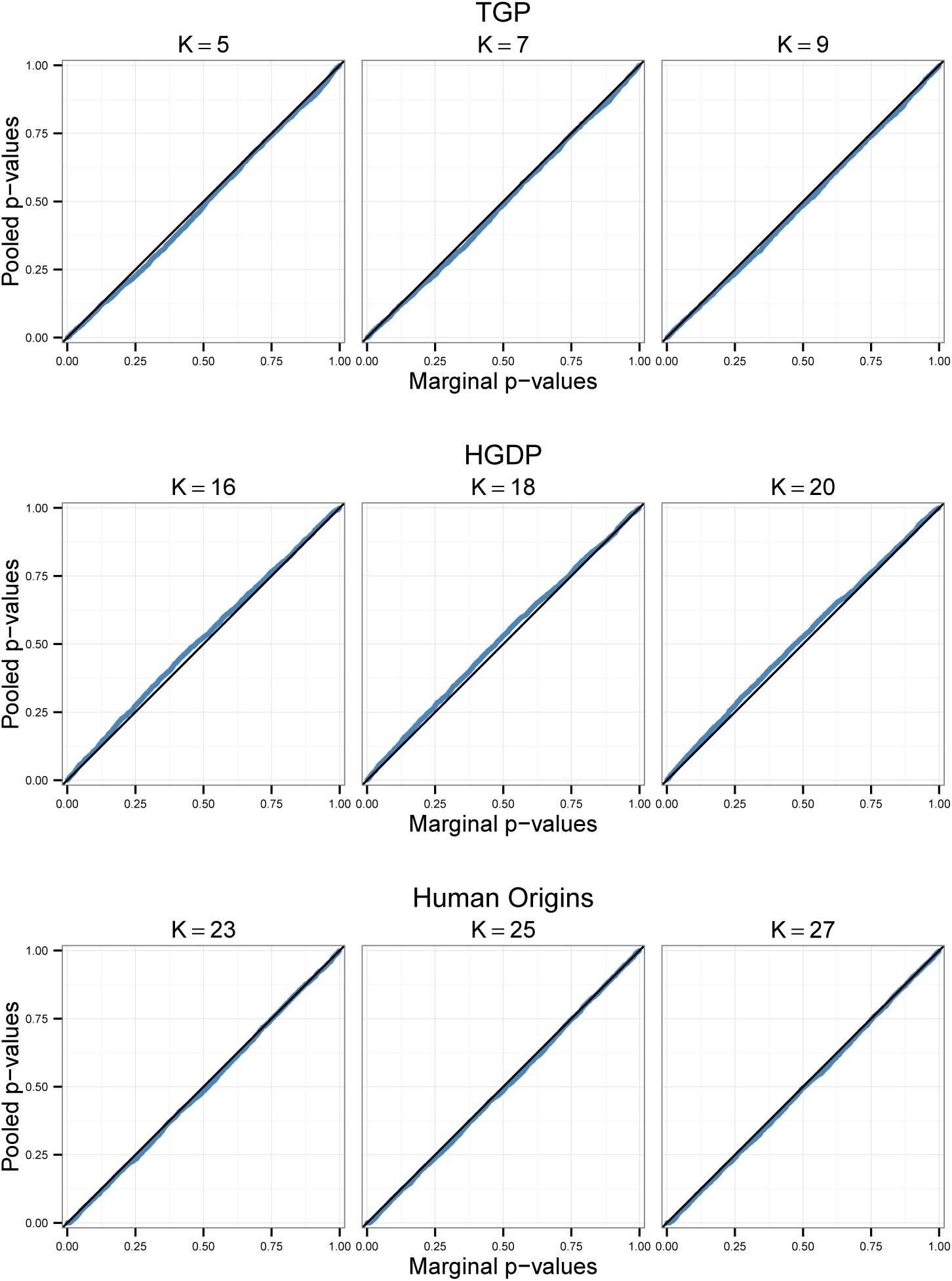
Quantile-quantile plots comparing p-values calculated using a SNP-specific marginal null distribution for each SNP versus the pooled null distribution for all three datasets. The quantile-quantile plots show a random sample of 5000 SNPs for each dataset to reduce clutter. The marginal null distribution was formed by simulating a new null dataset for each sample from the null distribution (10,000 simulated datasets total). The pooled null distribution was formed by pooling across all SNPs for 3 simulated null datasets.

**Figure S2:**
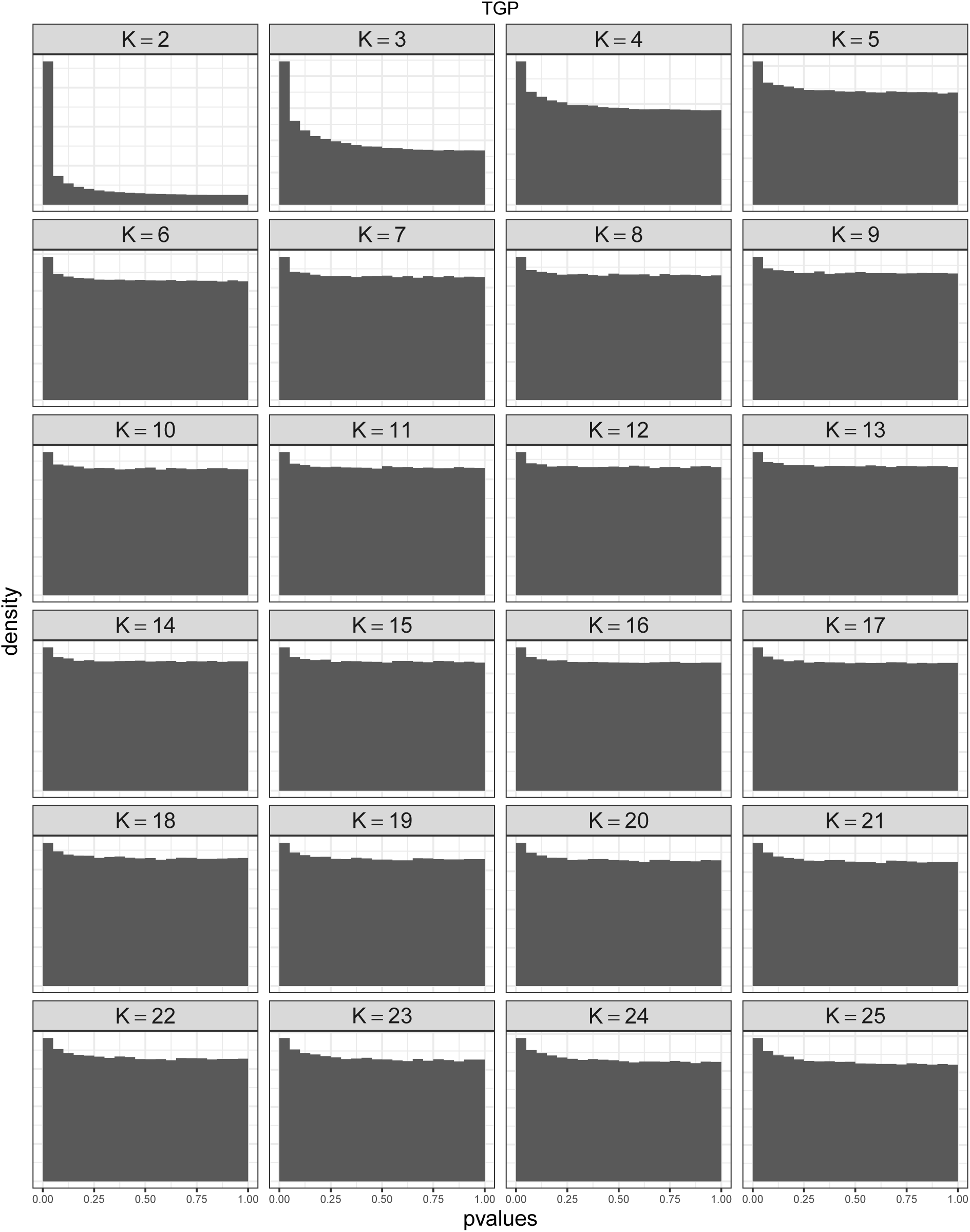
sHWE p-value histograms over a range of *K* for the TGP dataset. Note that for low *K*, population structure is only partially corrected for, resulting in a p-value distribution skewed towards zero. As *K* increases, skew towards zero reduces and the upper tail of the histogram becomes more Uniform(0,1), indicating that population structure is modeled more accurately resulting in minimal departures from sHWE.

**Figure S3:**
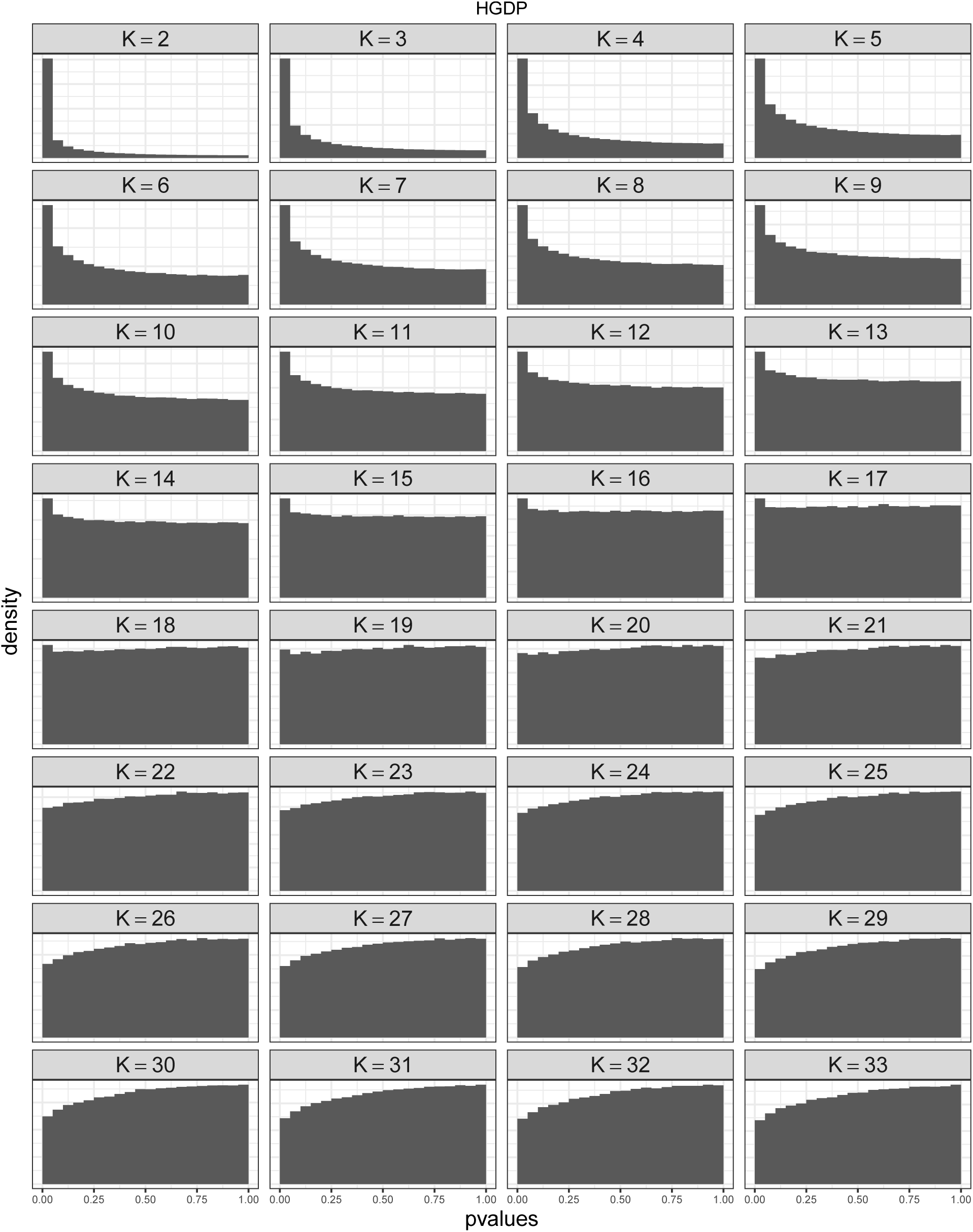
sHWE p-value histograms over a range of *K* for the HGDP dataset. Note that for low *K*, population structure is only partially corrected for, resulting in a p-value distribution skewed towards zero. As *K* increases, skew towards zero reduces and the upper tail of the histogram becomes more uniform, indicating that population structure is being better modeled.

**Figure S4:**
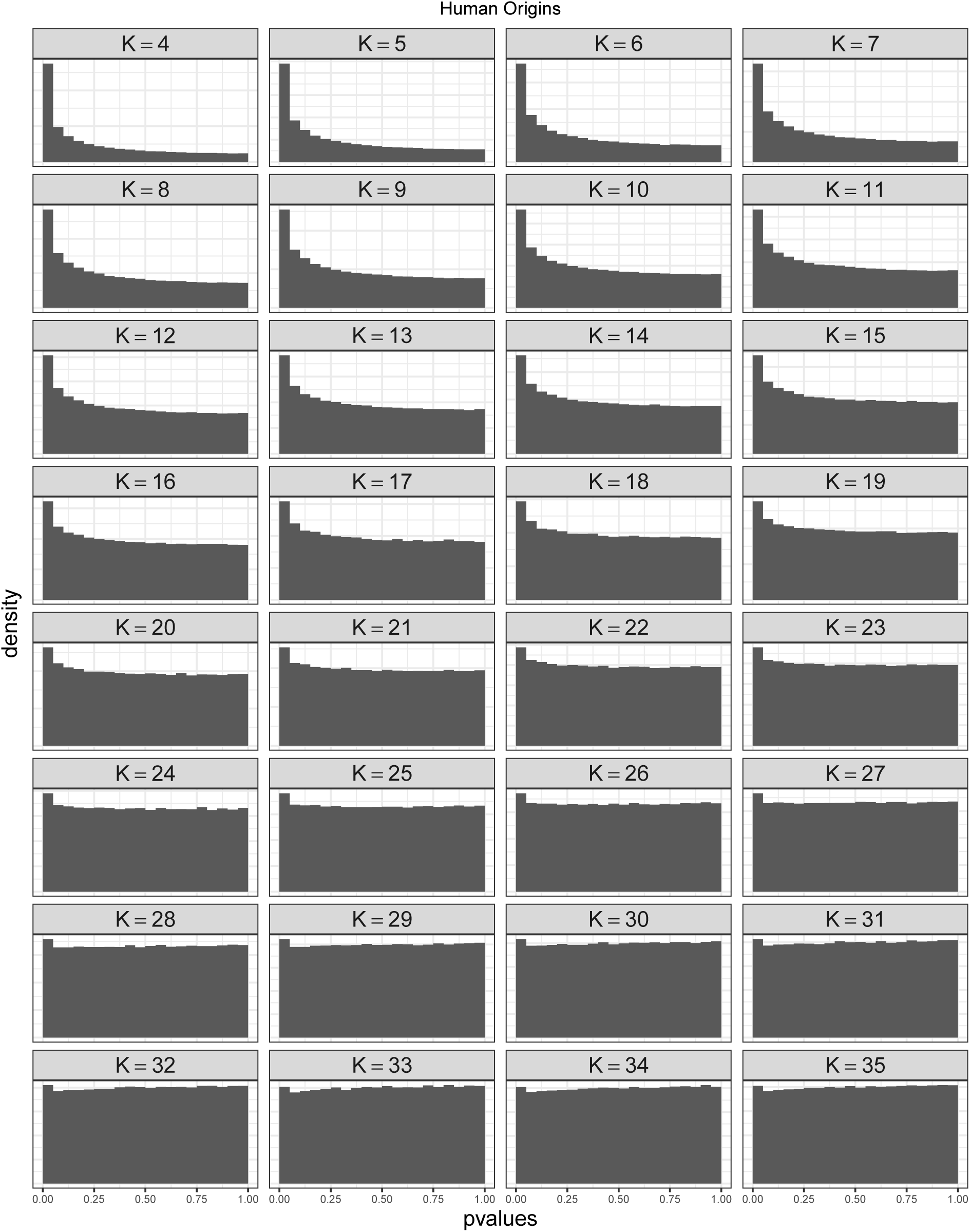
sHWE p-value histograms over a range of *K* for the Human Origins dataset. Note that for low *K*, population structure is only partially corrected for, resulting in a p-value distribution skewed towards zero. As *K* increases, skew towards zero reduces and the upper tail of the histogram becomes more uniform, indicating that population structure is being better modeled.

**Figure S5:**
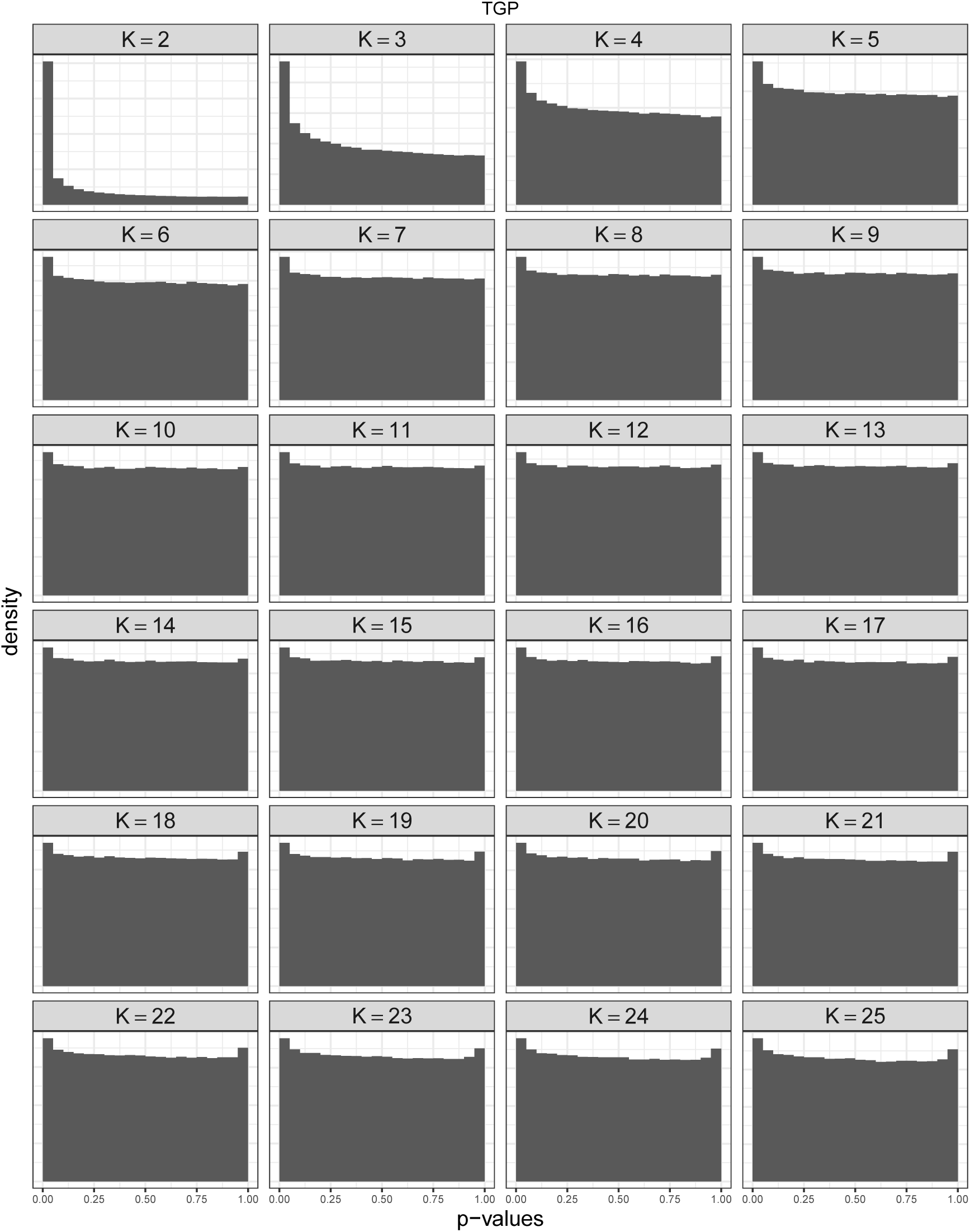
Structural HWE p-values for the TGP dataset using the truncated PCA approach to estimate allele frequencies. This is analogous to Figure S2.

**Figure S6:**
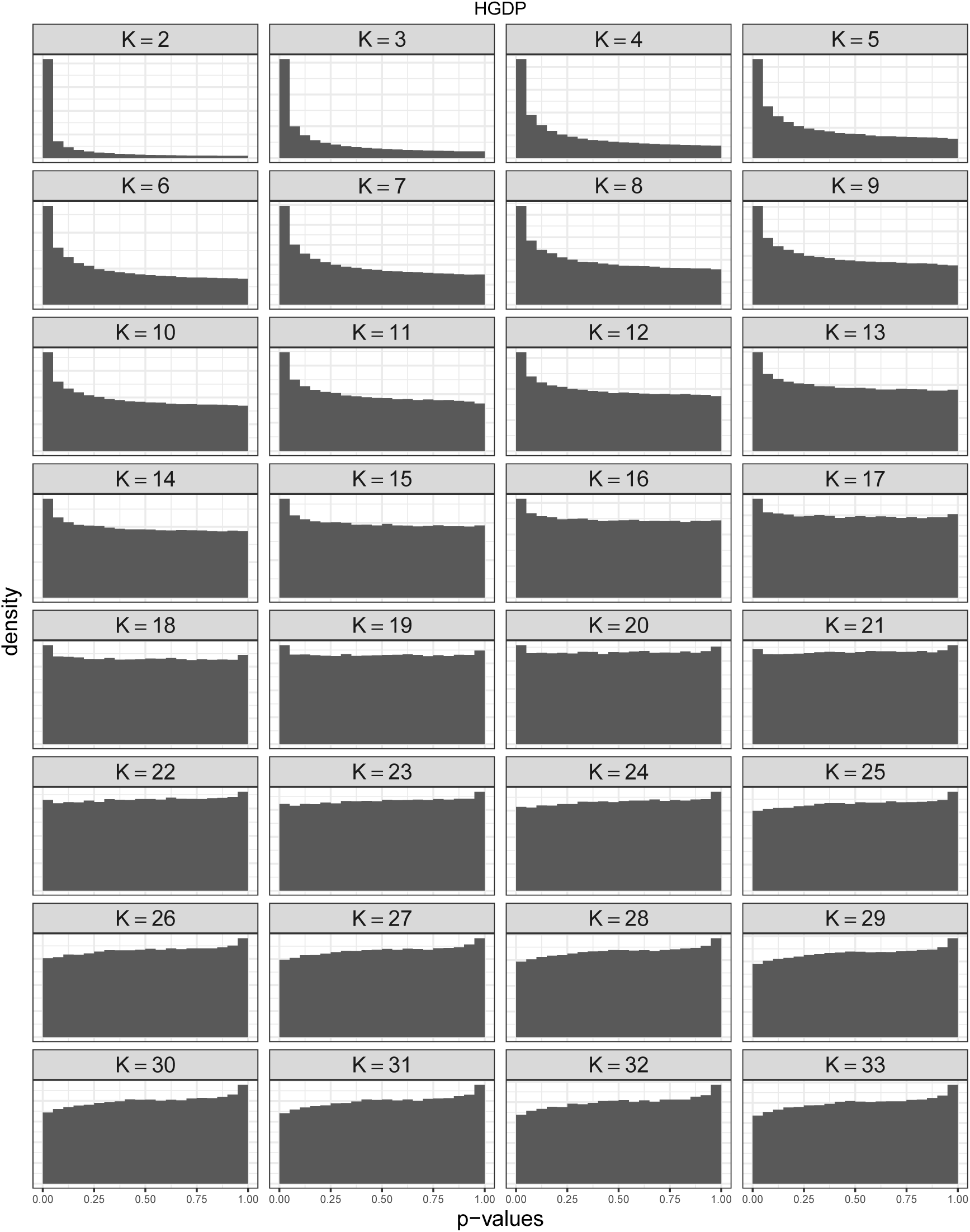
Structural HWE p-values for the HGDP dataset using the truncated PCA approach to estimate allele frequencies. This is analogous to Figure S3.

**Figure S7:**
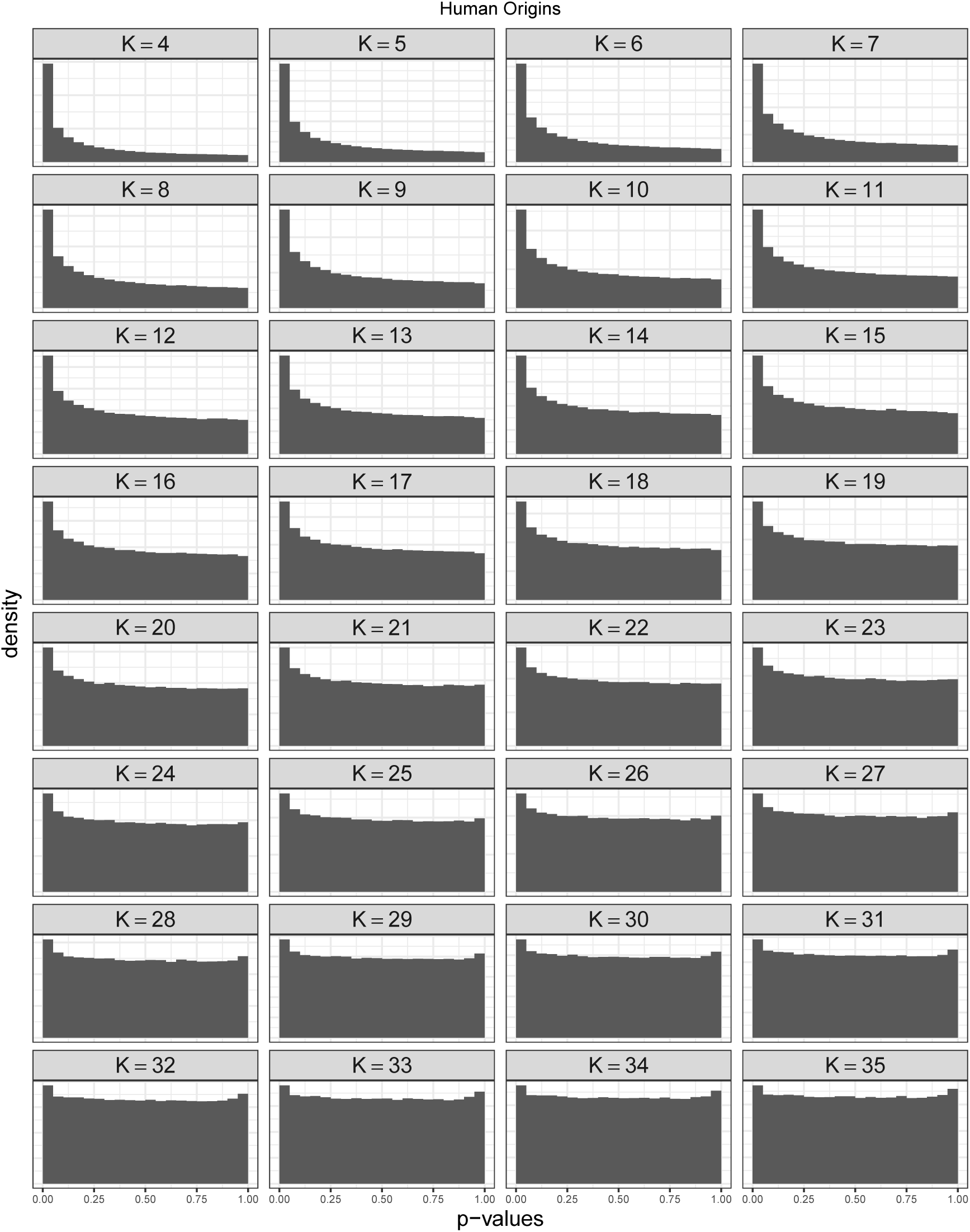
Structural HWE p-values for the HO dataset using the truncated PCA approach to estimate allele frequencies. This is analogous to Figure S4.

**Figure S8:**
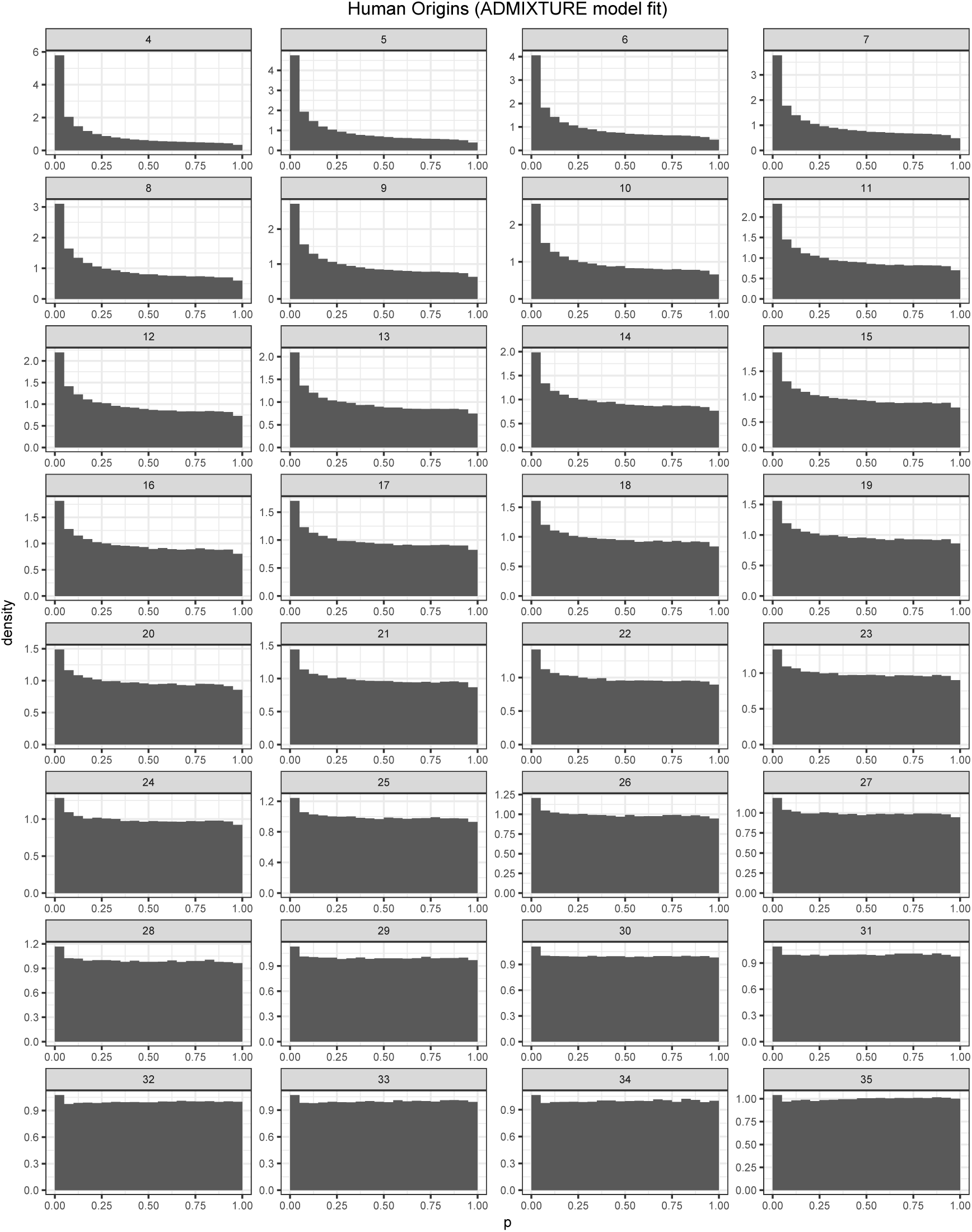
Structural HWE p-values for the HO dataset using the ADMIXTURE method to estimate allele frequencies. This is analogous to Figure S4.

**Figure S9:**
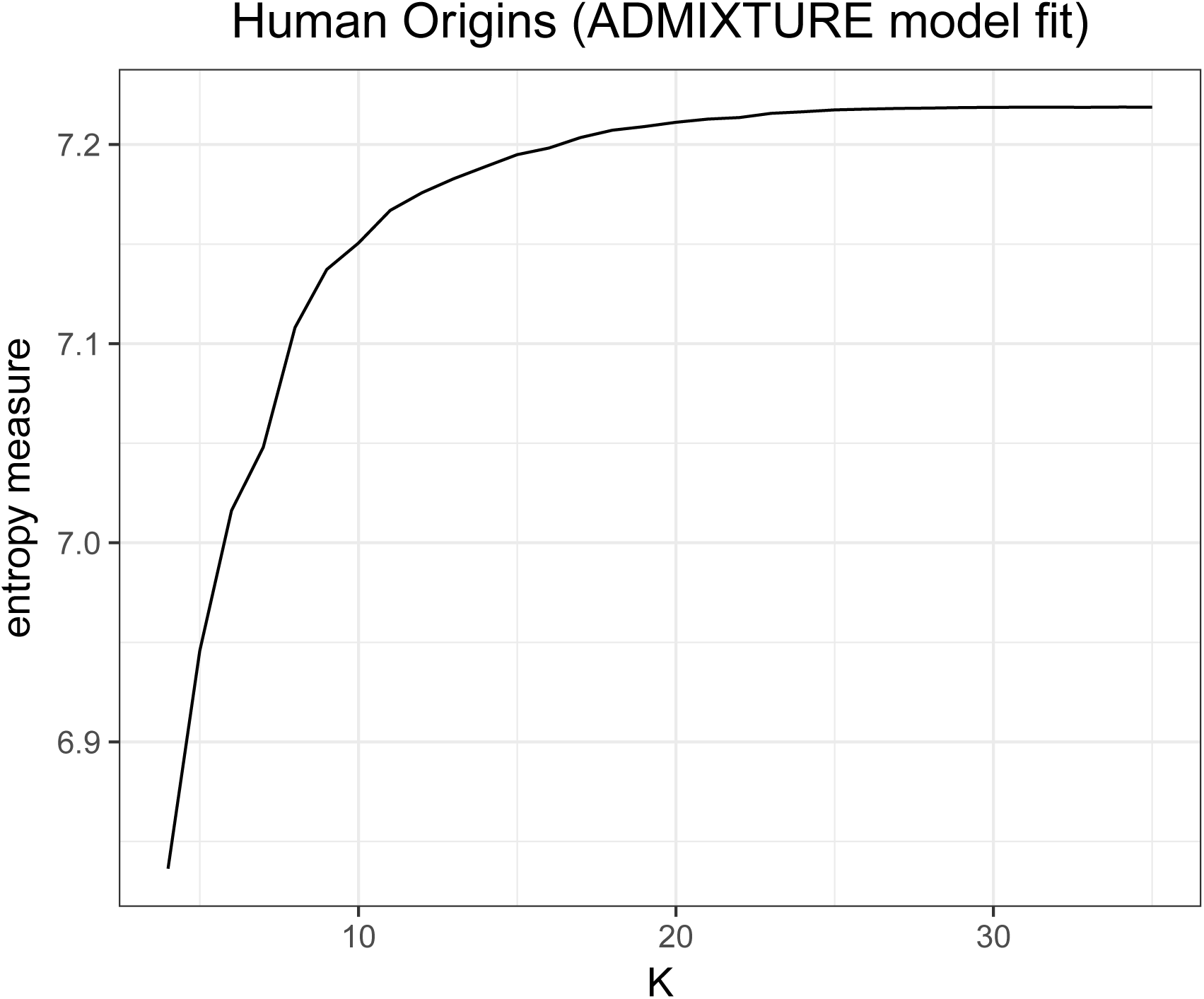
Entropy measure over a range of *K* for the HO dataset using the ADMIXTURE method to estimate allele frequencies.

**Figure S10:**
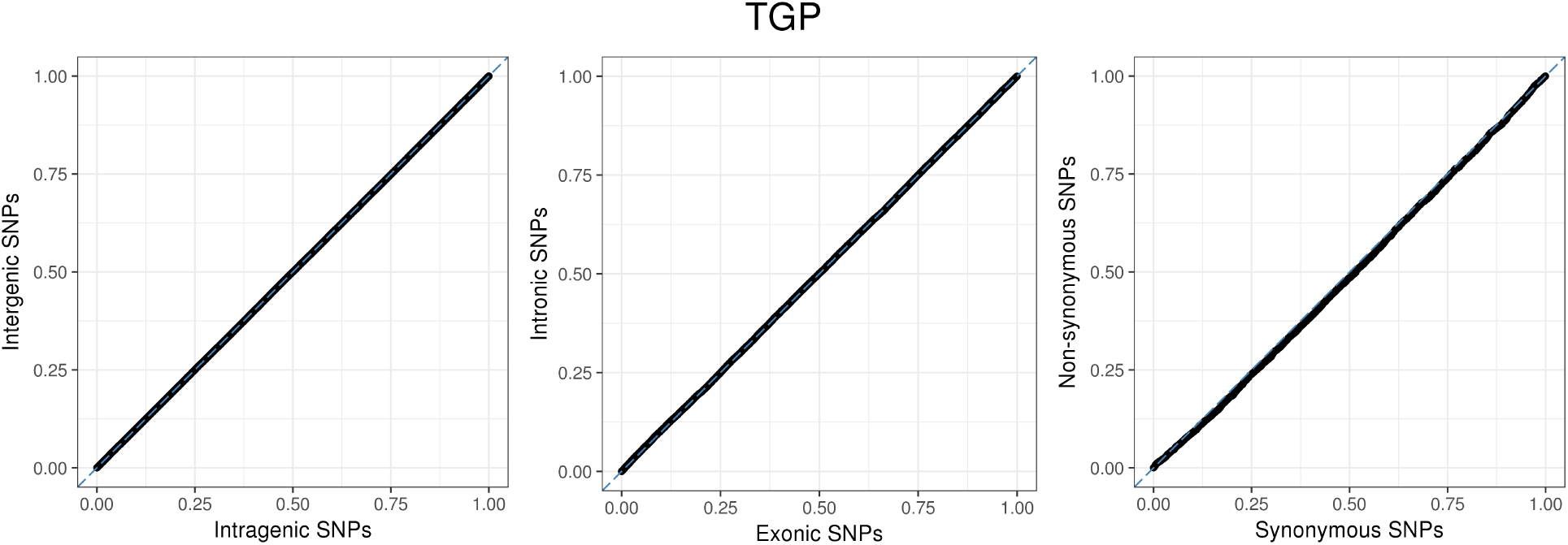
Quantile-quantile plots of structural HWE p-values for the TGP dataset at *K* = 12 comparing successively more fine-scale functional dichotomies for SNPs. The coarsest categories are intergenic SNPs versus intragenic SNPs. Then, among the intragenic SNPs, the comparison is between intronic SNPs versus exonic SNPs. Lastly, among the exonic SNPs, the comparison is between non-synonymous SNPs versus synonymous SNPs. Functional annotations are based on dbSNP build 147.

**Figure S11:**
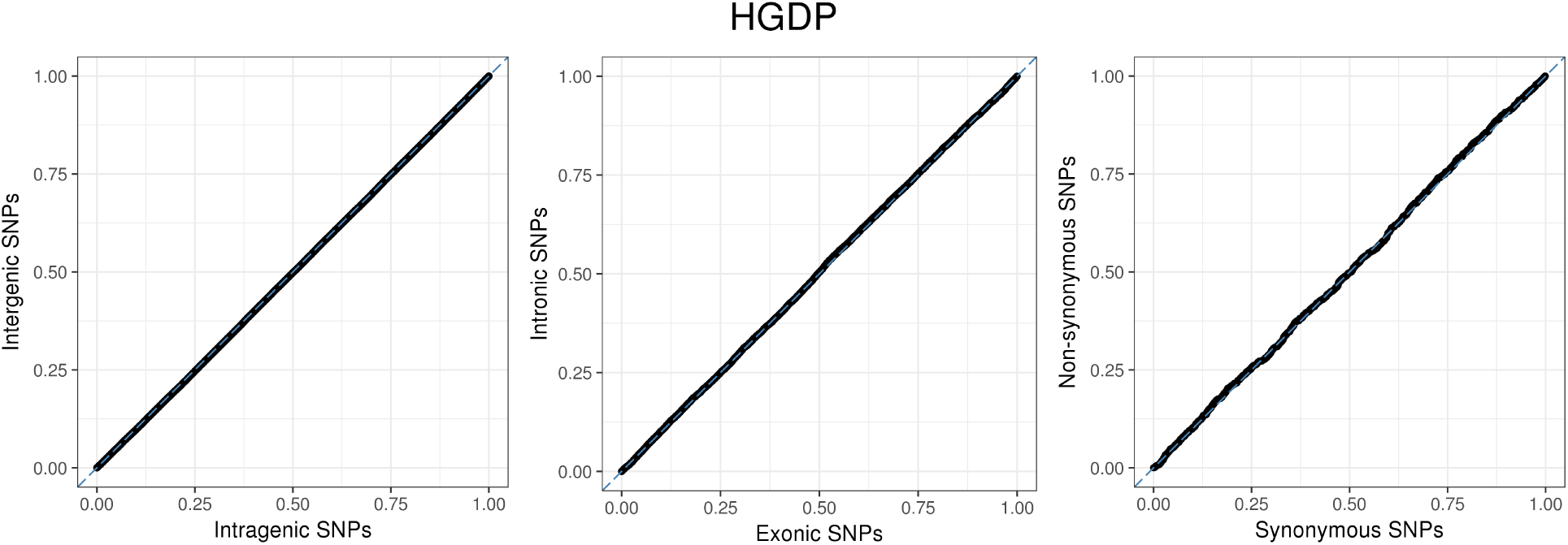
Quantile-quantile plots of structural HWE p-values for the HGDP dataset at *K* = 16 comparing successively more fine-scale functional dichotomies for SNPs. The coarsest categories are intergenic SNPs versus intragenic SNPs. Then, among the intragenic SNPs, the comparison is between intronic SNPs versus exonic SNPs. Lastly, among the exonic SNPs, the comparison is between non-synonymous SNPs versus synonymous SNPs. Functional annotations are based on dbSNP build 147.

**Figure S12:**
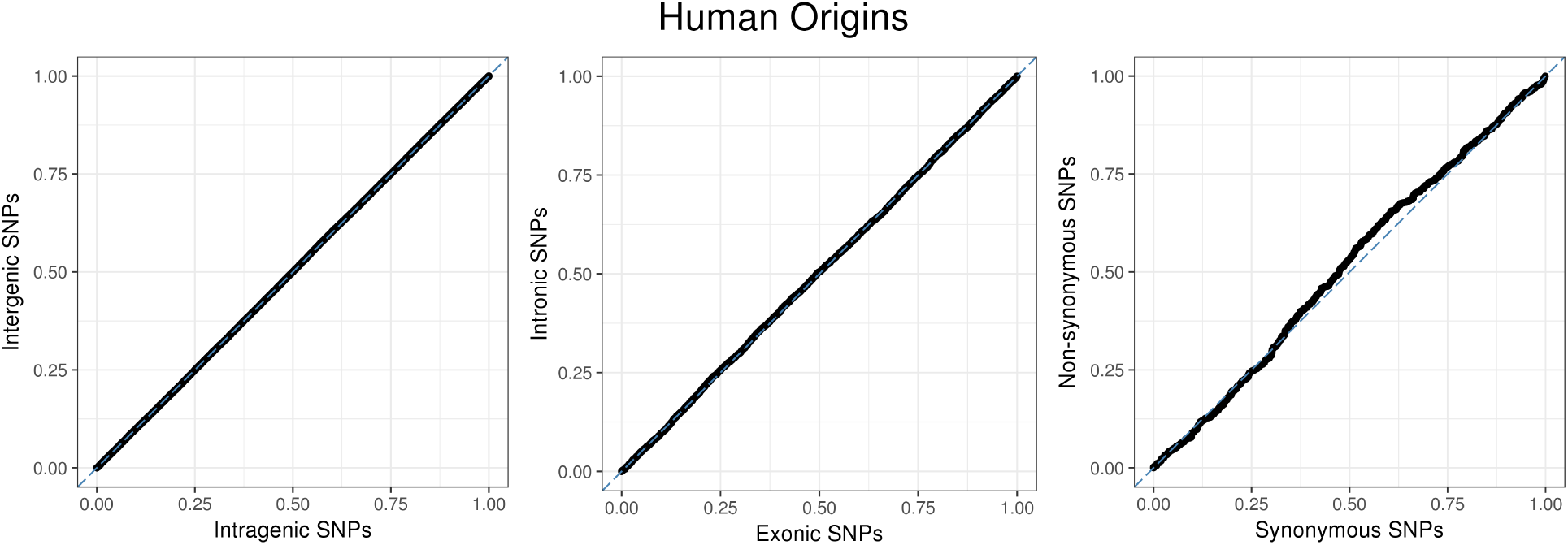
Quantile-quantile plots of structural HWE p-values for the Human Origins dataset at *K* = 25 comparing successively more fine-scale functional dichotomies for SNPs. The coarsest categories are inter-genic SNPs versus intragenic SNPs. Then, among the intragenic SNPs, the comparison is between intronic SNPs versus exonic SNPs. Lastly, among the exonic SNPs, the comparison is between non-synonymous SNPs versus synonymous SNPs. Functional annotations are based on dbSNP build 147.

**Figure S13:**
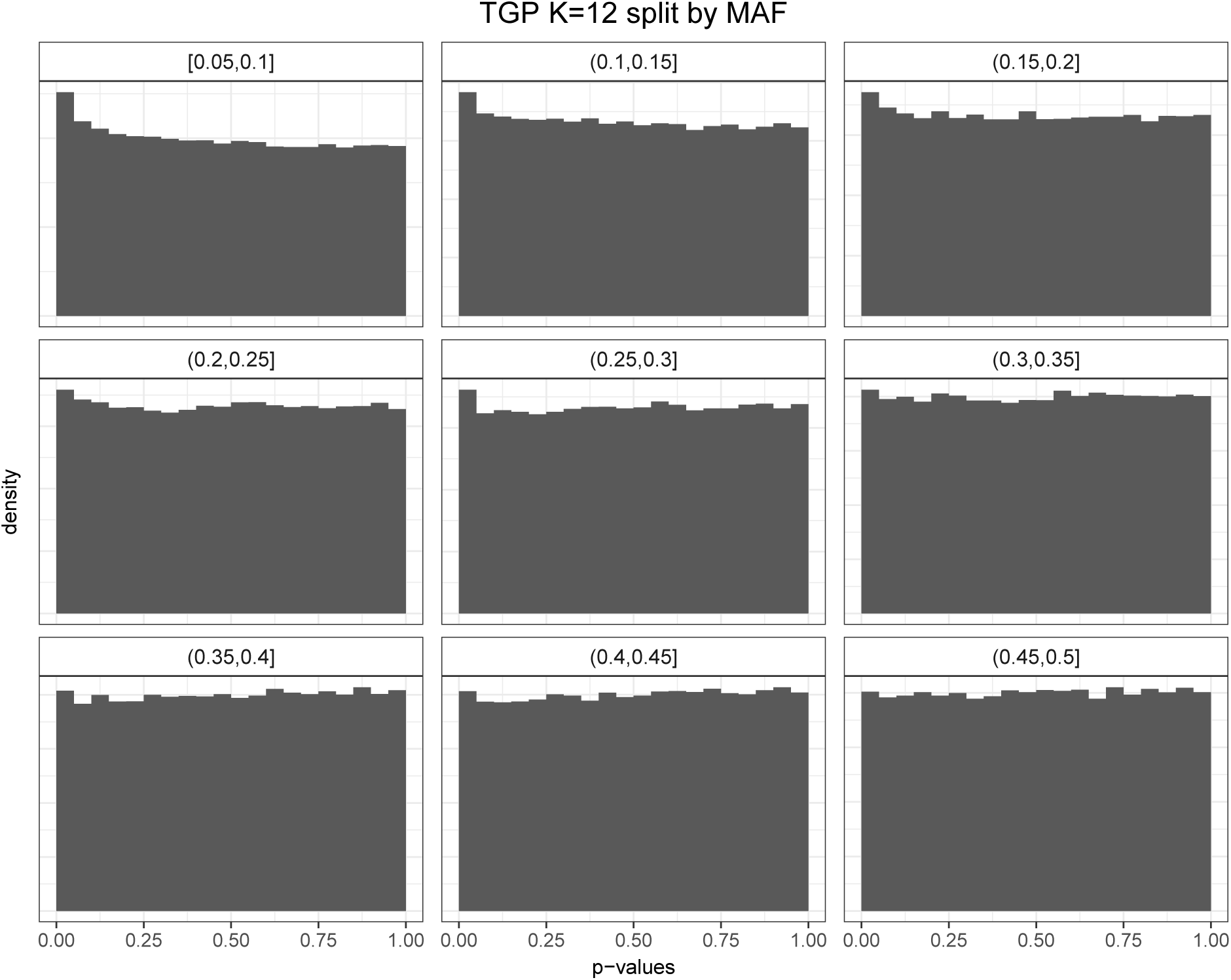
Structural HWE p-value histograms split by minor allele frequency for the TGP dataset (*K* = 12). There is minimal difference between bins.

**Figure S14:**
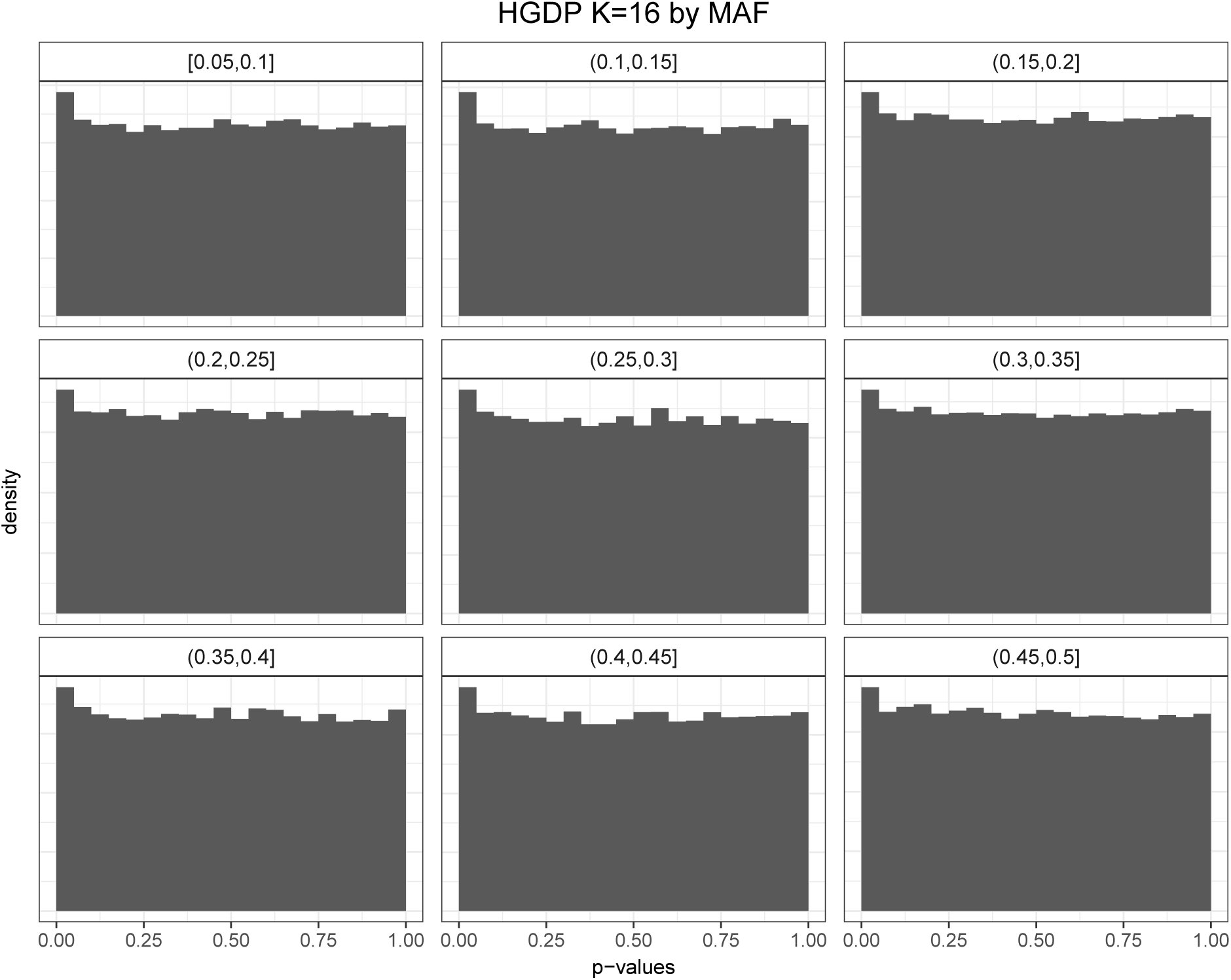
Structural HWE p-value histograms split by minor allele frequency for the HGDP dataset (K = 16). There is minimal difference between bins.

**Figure S15:**
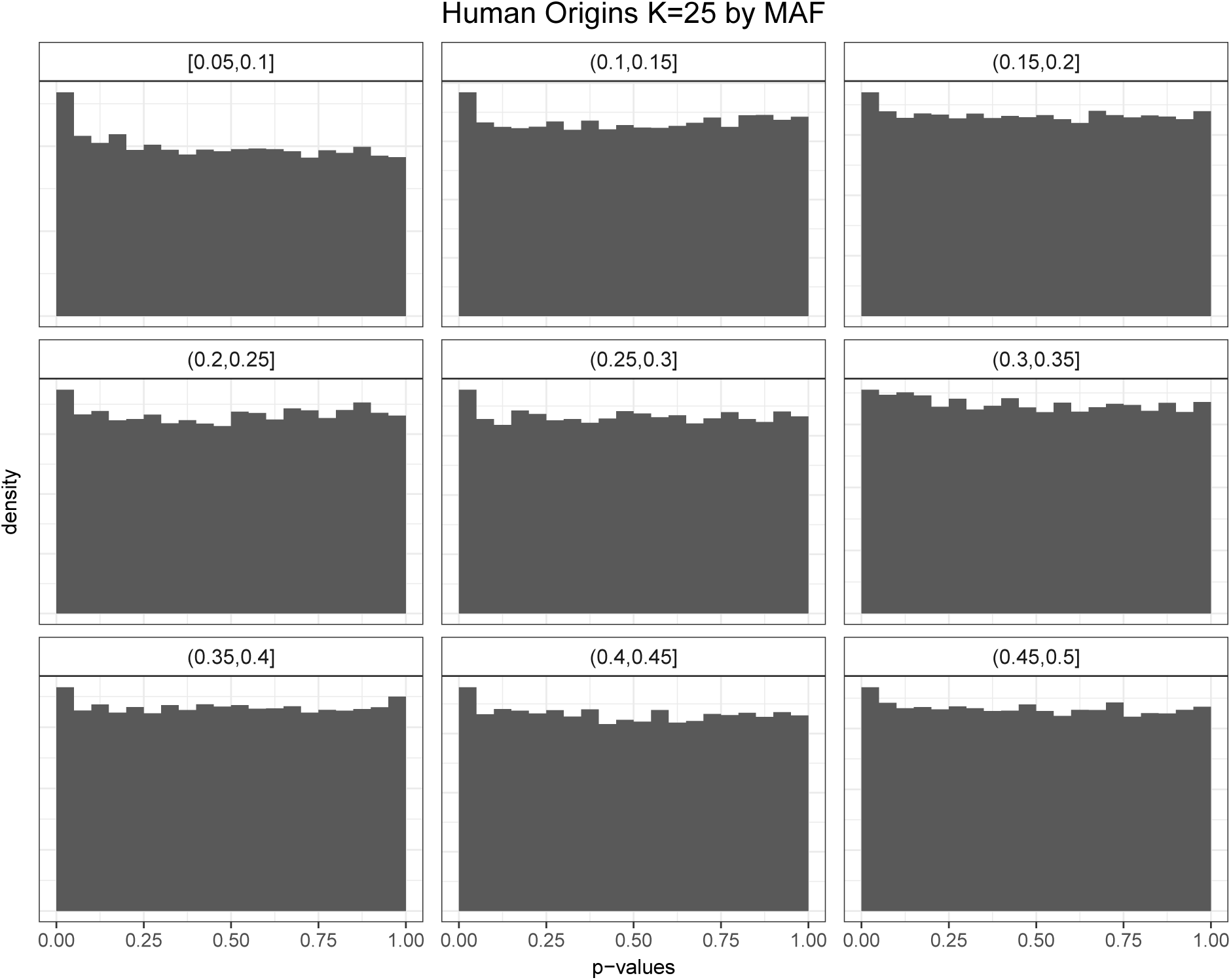
Structural HWE p-value histograms split by minor allele frequency for the Human Origins dataset (*K* = 25). There is minimal difference between bins.

**Figure S16:**
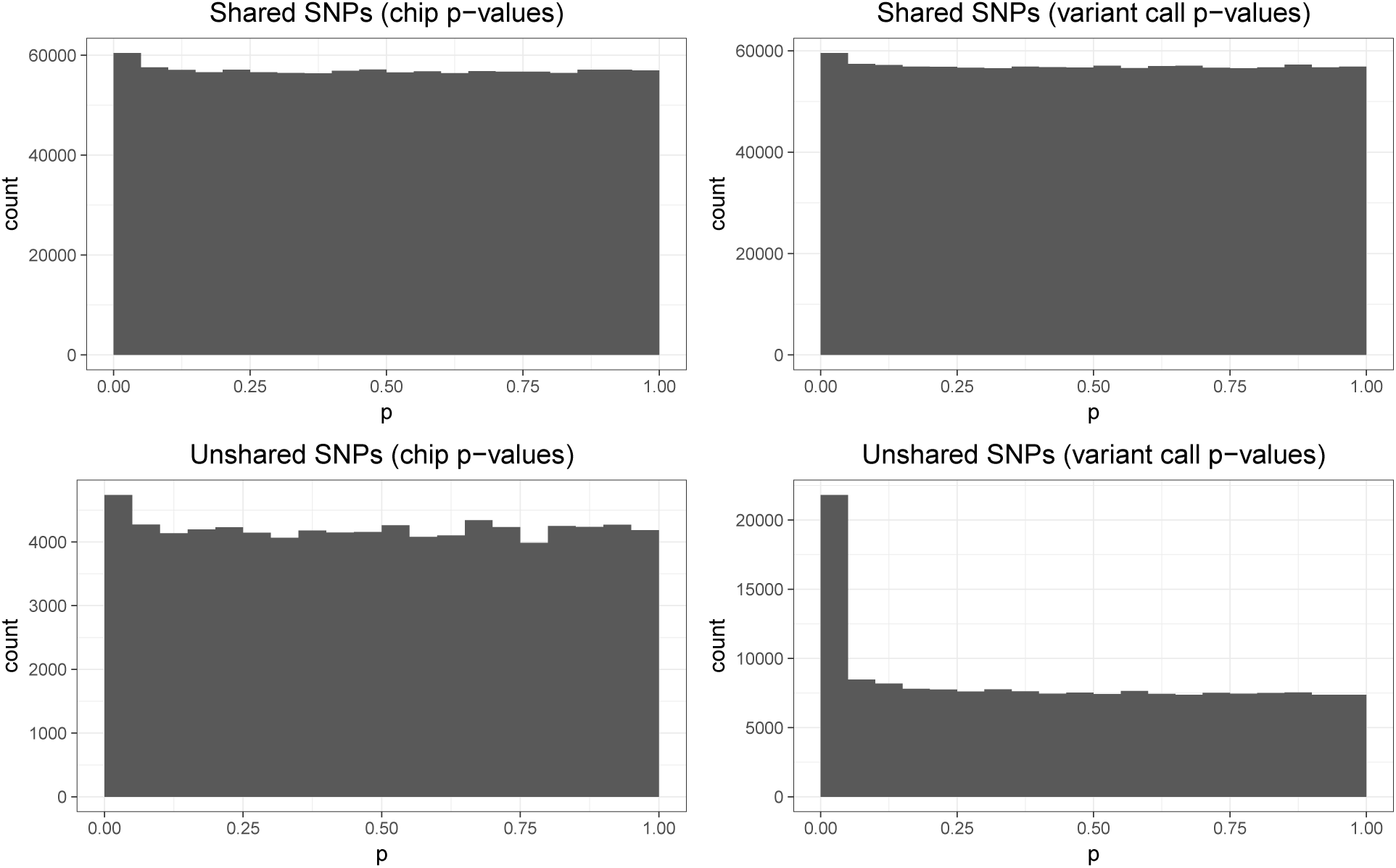
Histogram of sHWE p-values for the TGP chip and variant call datasets, using all SNPs and split between shared and unshared SNPs. The shared SNPs have similar distributions which shows that their results are conserved, while the unshared SNPs show that there are many more SNPs in the integrated callset which are significant.

## References

[1] Winkler, T. W., Day, F. R., Croteau-Chonka, D. C., Wood, A. R., Locke, A. E., Mägi, R., Ferreira, T., Fall, T., Graff, M., Justice, A. E., Luan, J., Gustafsson, S., Randall, J. C., Vedantam, S., Workalemahu, T., Kilpeläinen, T. O., Scherag, A., Esko, T., Kutalik, Z., Heid, I. M., and Loos, R. J. F. Quality control and conduct of genome-wide association meta-analyses. Nat. Protoc. 9(5) (2014).

[2] Anderson, C. A., Pettersson, F. H., Clarke, G. M., Cardon, L. R., Morris, A. P., and Zondervan, K. T. Data quality control in genetic case-control association studies. Nat.Protoc. 5(9) (2010).

[3] Pritchard, J. K., Stephens, M., and Donnelly, P. Inference of population structure using multilocus genotype data. Genetics 155(2), 945–59, jun (2000).

[4] Patterson, N., Price, A. L., and Reich, D. Population structure and eigenanalysis. PLoS Genet. 2(12), e190 (2006).

[5] Yang, J., Lee, S. H., Goddard, M. E., and Visscher, P. M. GCTA: A tool for genome-wide complex trait analysis. Am. J. Hum. Genet. 88(1), 76–82 (2011).

[6] Price, A. L., Patterson, N. J., Plenge, R. M., Weinblatt, M. E., Shadick, N. a., and Reich, D. Principal components analysis corrects for stratification in genome-wide association studies. Nat. Genet. 38(8), 904–9 (2006).

[7] Li, J. Z., Absher, D. M., Tang, H., Southwick, A. M., Casto, A. M., Ramachandran, S., Cann, H. M., Barsh, G. S., Feldman, M., Cavalli-Sforza, L. L., and Myers, R. M. Worldwide human relationships inferred from genome-wide patterns of variation. Science 319(5866), 1100–4 (2008).

[8] Coop, G., Pickrell, J. K., Novembre, J., Kudaravalli, S., Li, J., Absher, D., Myers, R. M., Cavalli-Sforza, L. L., Feldman, M. W., and Pritchard, J. K. The role of geography in human adaptation. PLoS Genet. 5(6) (2009).

[9] Gormley, P., Anttila, V., Winsvold, B. S., Palta, P., Esko, T., Pers, T. H., Farh, K.-H., Cuenca-Leon, E., Muona, M., Furlotte, N. A., Kurth, T., Ingason, A., McMahon, G., Ligthart, L., Terwindt, G. M., Kallela, M., Freilinger, T. M., Ran, C., Gordon, S. G., Stam, A. H., Steinberg, S., Borck, G., Koiranen, M., Quaye, L., Adams, H. H. H., Lehtimäki, T., Sarin, A.-P., Wedenoja, J., Hinds, D. A., Buring, J. E., Schürks, M., Ridker, P. M., Hrafnsdottir, M. G., Stefansson, H., Ring, S. M., Hottenga, J.-J., Penninx, B. W. J. H., Färkkilä, M., Artto, V., Kaunisto, M., Vepsäläinen, S., Malik, R., Heath, A. C., Madden, P. A. F., Martin, N. G., Montgomery, G. W., Kurki, M. I., Kals, M., Mägi, R., Pärn, K., Hämäläinen, E., Huang, H., Byrnes, A. E., Franke, L., Huang, J., Stergiakouli, E., Lee, P. H., Sandor, C., Webber, C., Cader, Z., Muller-Myhsok, B., Schreiber, S., Meitinger, T., Eriksson, J. G., Salomaa, V., Heikkilä, K., Loehrer, E., Uitterlinden, A. G., Hofman, A., van Duijn, C. M., Cherkas, L., Pedersen, L. M., Stubhaug, A., Nielsen, C. S., Männikkö, M., Mihailov, E., Milani, L., Göbel, H., Esserlind, A.-L., Christensen, A. F., Hansen, T. F., Werge, T., Anttila, V., Artto, V., Belin, A. C., Boomsma, D. I., Børte, S., Chasman, D. I., Cherkas, L., Christensen, A. F., Cormand, B., Cuenca-Leon, E., Smith, G. D., Dichgans, M., van Duijn, C., Eising, E., Esko, T., Esserlind, A.-L., Ferrari, M., Frants, R. R., Freilinger, T. M., Furlotte, N. A., Gormley, P., Griffiths, L., Hamalainen, E., Hansen, T. F., Hiekkala, M., Ikram, M. A., Ingason, A., Järvelin, M.-R., Kajanne, R., Kallela, M., Kaprio, J., Kaunisto, M., Kubisch, C., Kurki, M., Kurth, T., Launer, L., Lehtimaki, T., Lessel, D., Ligthart, L., Litterman, N., van den Maagdenberg, A. M. J. M., Macaya, A., Malik, R., Mangino, M., McMahon, G., Muller-Myhsok, B., Neale, B. M., Northover, C., Nyholt, D. R., Olesen, J., Palotie, A., Palta, P., Pedersen, L. M., Pedersen, N., Posthuma, D., Pozo-Rosich, P., Pressman, A., Quaye, L., Raitakari, O., Schürks, M., Sintas, C., Stefansson, K., Stefansson, H., Steinberg, S., Strachan, D., Terwindt, G. M., Vila-Pueyo, M., Wessman, M., Winsvold, B. S., Wrenthal, W., Zhao, H., Zwart, J.-A., Kaprio, J., Aromaa, A. J., Raitakari, O., Ikram, M. A., Spector, T., Järvelin, M.-R., Metspalu, A., Kubisch, C., Strachan, D. P., Ferrari, M. D., Belin, A. C., Dichgans, M., Wessman, M., van den Maagdenberg, A. M. J. M., Zwart, J.-A., Boomsma, D. I., Smith, G. D., Stefansson, K., Eriksson, N., Daly, M. J., Neale, B. M., Olesen, J., Chasman, D. I., Nyholt, D. R., and Palotie, A. Meta-analysis of 375,000 individuals identifies 38 susceptibility loci for migraine. Nat. Genet. 48(8) (2016).

[10] Song, M., Hao, W., and Storey, J. D. Testing for genetic associations in arbitrarily structured populations. Nat. Genet. 47(5), 550–554 (2015).

[11] The 1000 Genomes Project Consortium. A map of human genome variation from population-scale sequencing. Nature 467(7319), 1061–73 (2010).

[12] The 1000 Genomes Project Consortium. A global reference for human genetic variation. Nature 526(7571) (2015).

[13] Bryc, K., Velez, C., Karafet, T., Moreno-Estrada, A., Reynolds, A., Auton, A., Hammer, M., Bustamante, C. D., and Ostrer, H. Genome-wide patterns of population structure and admixture among Hispanic/Latino populations. Proc. Natl. Acad. Sci. 107(Supplement_2), 8954–8961, may (2010).

[14] Thornton, T., Tang, H., Hoffmann, T. J., Ochs-Balcom, H. M., Caan, B. J., and Risch, N. Estimating Kinship in Admixed Populations. Am. J. Hum. Genet. 91(1), 122–138, jul (2012).

[15] Moreno-Estrada, A., Gignoux, C. R., Fernandez-Lopez, J. C., Zakharia, F., Sikora, M., Contreras, A. V., Acuna-Alonzo, V., Sandoval, K., Eng, C., Romero-Hidalgo, S., Ortiz-Tello, P., Robles, V., Kenny, E. E., Nuno-Arana, I., Barquera-Lozano, R., Macin-Perez, G., Granados-Arriola, J., Huntsman, S., Galanter, J. M., Via, M., Ford, J. G., Chapela, R., Rodriguez-Cintron, W., Rodriguez-Santana, J. R., Romieu, I., Sienra-Monge, J. J., Navarro, B. d. R., London, S. J., Ruiz-Linares, A., Garcia-Herrera, R., Estrada, K., Hidalgo-Miranda, A., Jimenez-Sanchez, G., Carnevale, A., Soberon, X., Canizales-Quinteros, S., Rangel-Villalobos, H., Silva-Zolezzi, I., Burchard, E. G., and Bustamante, C. D. The genetics of Mexico recapitulates Native American substructure and affects biomedical traits. Science (80-.). 344(6189), 1280–1285, jun (2014).

[16] Hao, W., Song, M., and Storey, J. D. Probabilistic models of genetic variation in structured populations applied to global human studies. Bioinformatics 32(5) (2016).

[17] Cann, H. M., de Toma, C., Cazes, L., and Legrand, M. F. A human genome diversity cell line panel. Science (80-.). 296(5566), 261 (2002).

[18] Lazaridis, I., Patterson, N., Mittnik, A., Renaud, G., Mallick, S., Sudmant, P. H., Schraiber, J. G., Castellano, S., Kirsanow, K., Economou, C., Bollongino, R., Fu, Q., Bos, K., Nordenfelt, S., de Filippo, C., Prüfer, K., Sawyer, S., Posth, C., Haak, W., Hallgren, F., Fornander, E., Ayodo, G., Babiker, H. a., Balanovska, E., Balanovsky, O., Ben-Ami, H., Bene, J., Berrada, F., Brisighelli, F., Busby, G. B., Cali, F., Churnosov, M., Cole, D. E., Damba, L., Delsate, D., van Driem, G., Dryomov, S., Fedorova, S. a., Francken, M., Gallego Romero, I., Gubina, M., Guinet, J.-M., Hammer, M., Henn, B., Helvig, T., Hodoglugil, U., Jha, A. R., Kittles, R., Khusnutdinova, E., Kivisild, T., Kučinskas, V., Khusainova, R., Kushniarevich, A., Laredj, L., Litvinov, S., Mahley, R. W., Melegh, B., Metspalu, E., Mountain, J., Nyambo, T., Osipova, L., Parik, J., Platonov, F., Posukh, O. L., Romano, V., Rudan, I., Ruizbakiev, R., Sahakyan, H., Salas, A., Starikovskaya, E. B., Tarekegn, A., Toncheva, D., Turdikulova, S., Uktveryte, I., Utevska, O., Voevoda, M., Wahl, J., Zalloua, P., Yepiskoposyan, L., Zemunik, T., Cooper, A., Capelli, C., Thomas, M. G., Tishkoff, S. a., Singh, L., Thangaraj, K., Villems, R., Comas, D., Sukernik, R., Metspalu, M., Meyer, M., Eichler, E. E., Burger, J., Slatkin, M., Pääbo, S., Kelso, J., Reich, D., and Krause, J. Ancient human genomes suggest three ancestral populations for present-day Europeans. Nature 513(7518), 409–413 (2014).

[19] Rosenberg, N. a. Standardized subsets of the HGDP-CEPH Human Genome Diversity Cell Line Panel, accounting for atypical and duplicated samples and pairs of close relatives. Ann. Hum. Genet. 70(Pt 6), 841–7 (2006).

[20] Wigginton, J. E., Cutler, D. J., and Abecasis, G. R. A note on exact tests of Hardy-Weinberg equilibrium. Am. J. Hum. Genet. 76(5), 887–893 (2005).

[21] Alexander, D. H., Novembre, J., and Lange, K. Fast model-based estimation of ancestry in unrelated individuals. Genome Res. 19, 1655–1664 (2009).

[22] Wasser, S. K., Shedlock, A. M., Comstock, K., Ostrander, E. A., Mutayoba, B., and Stephens, M. Assigning African elephant DNA to geographic region of origin: Applications to the ivory trade. Proc. Natl. Acad. Sci. 101(41), 14847–14852, oct (2004).

[23] Corander, J., Sirén, J., and Arjas, E. Bayesian spatial modeling of genetic population structure. Comput. Stat. 23(1), 111–129, jan (2008).

[24] Chung, N. C. and Storey, J. D. Statistical significance of variables driving systematic variation. Bioinformatics 31(4), 545–554, aug (2013).

[25] Efron, B. and Tibshirani, R. An introduction to the bootstrap. Chapman & Hall/CRC, (1993).

[26] Leek, J. T. and Storey, J. D. The Joint Null Criterion for Multiple Hypothesis Tests. Stat. Appl. Genet. Mol. Biol. 10(1), 1–22 (2011).

[27] Gopalan, P., Hao, W., Blei, D. M., and Storey, J. D. Scaling probabilistic models of genetic variation to millions of humans. Nat. Genet. 48(12), 1587–1590, nov (2016).

[28] Storey, J. D. and Tibshirani, R. Statistical significance for genomewide studies. Proc. Natl. Acad. Sci. U. S. A. 100(16), 9440–9445 (2003).

[29] Hinds, D. A. Whole-Genome Patterns of Common DNA Variation in Three Human Populations. Science (80-.). 307(5712), 1072–1079, feb (2005).

